# Extracellular Respiration is a Latent Energy Metabolism in *Escherichia coli*

**DOI:** 10.1101/2024.05.30.596743

**Authors:** Biki Bapi Kundu, Jayanth Krishnan, Richard Szubin, Arjun Patel, Bernhard O. Palsson, Daniel C. Zielinski, Caroline M. Ajo-Franklin

## Abstract

Diverse microbial species utilize redox shuttles to exchange electrons with their environment through mediated extracellular electron transfer (EET). This process maintains redox homeostasis and supports anaerobic survival across diverse microbial communities. Although mediated EET has been extensively leveraged for bioelectrocatalysis and bioelectronics for decades, fundamental questions remain about how these redox shuttles are reduced within cells and their bioenergetic implications. This knowledge gap limits our understanding of the physiological roles of mediated EET in various microbes and hampers the development of efficient microbial electrochemical technologies. To address this, we developed a methodology integrating genome editing, electrochemistry, and systems biology to investigate the mechanism and bioenergetic implications of mediated EET in bacteria. Using this approach, we uncovered a mediated EET mechanism in *Escherichia coli*. In the absence of alternative electron sinks, the redox cycling of 2-hydroxy-1,4-naphthoquinone (HNQ) via a cytoplasmic nitroreductase enabled *E. coli* to respire and grow on an extracellular electrode. Genome-scale metabolic modeling suggested that HNQ- mediated EET offers a more energetically favorable route for supporting anaerobic growth than canonical fermentation. Transcriptome analysis revealed redox perturbations in response to HNQ and identified rapid metabolic adaptations that support growth. This work demonstrates that *E. coli* can grow independently of classical electron transport chains and fermentative pathways, unveiling a new type of anaerobic energy metabolism.

## INTRODUCTION

Electron flow drives energy metabolism, sustaining life at its core^1^. Energy metabolisms essentially transfer electrons to an electron sink through diverse respiratory and fermentative pathways. *Escherichia coli,* the cornerstone microbe for studying energy metabolism^2^, is conventionally known to reduce *intra*cellular electron sinks such as oxygen and nitrate^3,4^. Exoelectrogens, conversely, are specialized microbes that reduce extracellular electron sinks, such as metal (hydro)oxides, through a process called *extra*cellular electron transfer (EET)^5,6^. EET is pivotal in diverse environments, contributing to Earth’s biogeochemical cycles^7–9^ and microbial community dynamics in mammalian gastrointestinal tracts^10,11^. Moreover, EET pathways can connect electronics and living cells, enabling the electrical modulation of energy metabolism, gene expression, and overall cellular physiology^12–16^.

*E. coli is* conventionally regarded as a non-exoelectrogen and, because of its importance as a biotechnology workhorse, has been the focus of extensive engineering efforts to introduce EET^17–22^. However, the synthetically introduced EET registered only a modest electrical output, and unlike in model exoelectrogens, EET is uncoupled to growth^21,23–25^. Intriguingly, it has been known for over 20 years that *E. coli* can evolve to perform EET when interfaced with an extracellular electrode^26–35^. The reported EET levels from evolved *E. coli* are robust and comparable to pure cultures of model exoelectrogens^36^. The evolved EET is mediated by 2-hydroxy-1,4-naphthoquinone (HNQ), which is hypothesized to be biosynthesized and secreted by *E. coli*. These observations are of great interest because they suggest a latent exoelectrogenic mode of energy metabolism for anaerobic sustenance. Moreover, the ability of *E. coli* to perform HNQ-mediated EET coupled to growth could pave the way for constructing robust biosensors and synthetic metabolisms for carbon capture. Despite these opportunities, the mechanism and bioenergetic implications of HNQ-mediated EET remain unknown. This is because elucidating the precise mechanism of mediated EET requires probing numerous cellular enzymes that can reduce the redox shuttle, involving a trial-and-error approach with a substantial experimental burden^37,38^.

In this study, we addressed this gap by integrating high-fidelity genome editing, systems biology, and microbial electrochemistry to elucidate the mechanisms and implications of mediated EET in *E. coli*. With these tools, we dissected electron flow in *E. coli* metabolism, genetically altering the electron flux network and gauging the impact of the modified network on EET levels (Figure 1A). Using this approach, we found that a cytoplasmic nitroreductase NfsB selectively facilitates HNQ-mediated EET, efficiently driving catabolism by enabling NAD^+^ regeneration under resting conditions. Additionally, HNQ-mediated EET allows *E. coli* to respire on an anode and support unexpectedly robust anaerobic growth. Flux-balance analysis suggested that EET-mediated anaerobic metabolism offers substantial bioenergetic advantages over mixed-acid fermentation. Transcriptomic response to HNQ revealed substantial perturbation of redox pathways and a shift to support growth through anaplerotic catabolism of peptides. In summary, the discovered naphthoquinone-mediated EET pathway reveals a new type of anaerobic energy metabolism and lays the foundation for a minimal and generalizable method of engineering EET in microbes to improve planetary and human health.

**Figure 1:**
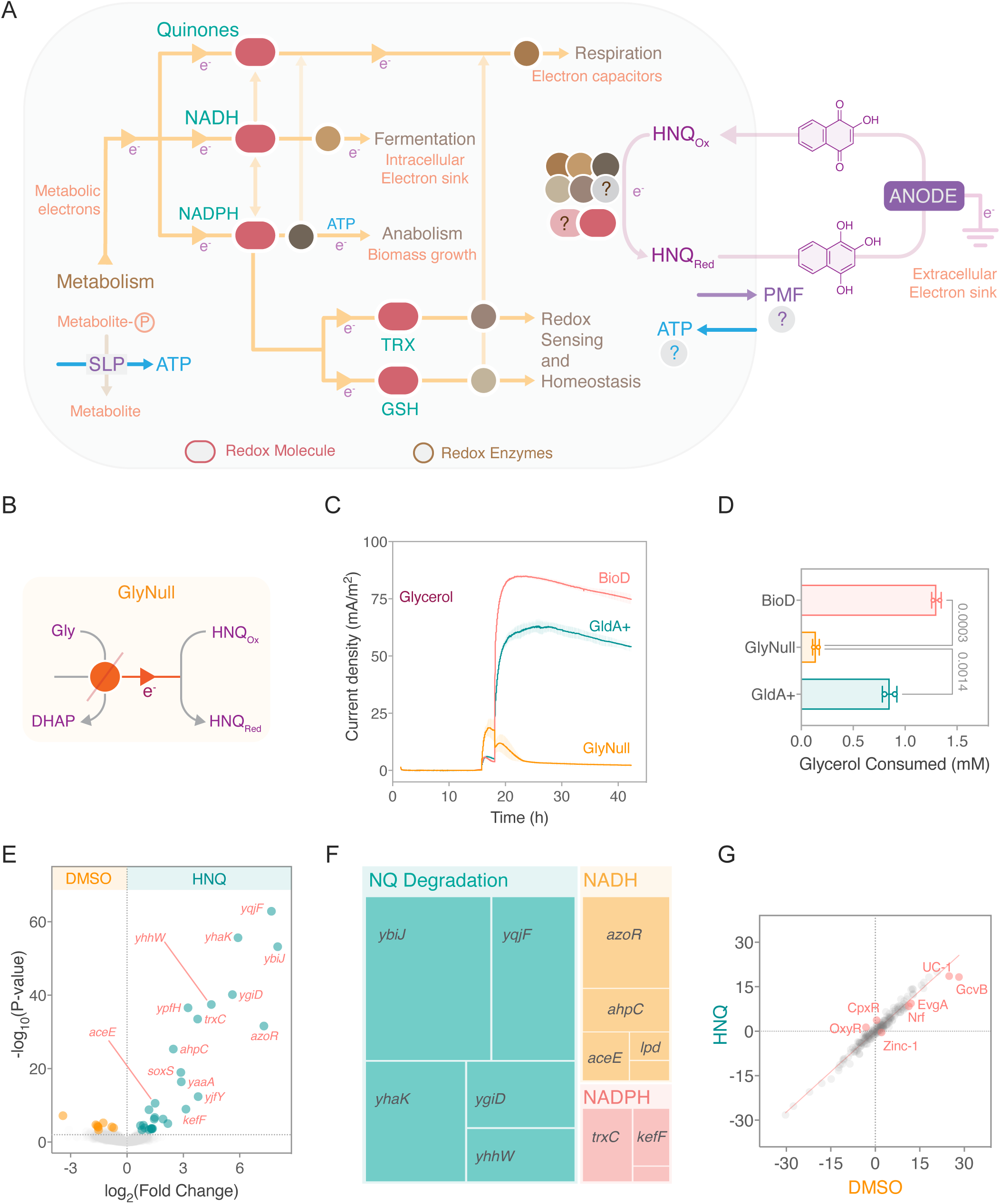
HNQ-mediated EET is coupled to catabolism. (A) A schematic illustrating that HNQ serves as a redox shuttle between cells and anode. Elucidating this EET mechanism is challenging because several cellular electron carriers, such as NADH, NADPH, inner membrane quinones, glutathione, and thioredoxin, can transfer electrons to HNQ either enzymatically or non-enzymatically. (B) The GlyNull strain was generated by deleting the glycerol consumption pathways in the BioD strain. (C) Chronoamperometry of BioD, GlyNull, and GldA^+^ strains under resting conditions, showing that only strains capable of consuming the available carbon source can perform EET. The strains in this experiment were supplemented with 20 mM glycerol in both pre-culture and biomass-generating cultures. (D) Only the BioD and GldA^+^ strains are capable of consuming glycerol, while GlyNull, lacking the glycerol oxidation pathways, does not. The data in (C) and (D) represent mean ± standard deviation obtained in duplicate BES experiments. The P-values are calculated using one-way ANOVA, with a significance threshold of 0.05. (E) Differential gene expression analysis (DEG) of BioD in DMSO alone and with HNQ under open circuit conditions, showing that the presence of HNQ upregulates a relatively small amount of gene expression. This data indicates that HNQ reduction is coupled with the induction of thioredoxin, *trxC* and the quinone reductase, *azoR*. Upregulation of *soxS* suggests that HNQ is an activator of the SoxRS, which defrays the toxic effects of electrophiles. (F) A Treemap of the explained variance of each DEG in the transcriptome of HNQ-treated cells, showing that differentially expressed genes are enriched with redox-associated genes. The evidence points to NQ degradation comprising the largest gene expression changes, followed by NADH, and NADPH associated gene sets. (G) Differential iModulon activity (DiMA) analysis of HNQ-treated BioD, showing that redox stress and metal stress gene sets are differentially regulated, along with the nrf operon. UC-1 is a complex iModulon containing prophage and redox genes. EvgA is associated with an acid and drug resistance two-component sensor. The data in (E), (F), and (G) represent mean ± standard deviation obtained in a triplicate no-EET BES setup, where the potentiostat was not connected to the BES. The BESs did not have anode, and cells were injected at the final OD_600_ of 0.25. The cells were treated with HNQ for three hours under resting conditions and harvested for RNA extraction.

## RESULTS

### The HNQ-mediated EET pathway depends on the oxidation of a carbon source

Based on prior studies, we hypothesized that exogenous HNQ could stimulate EET in *E. coli*^30,39^. To determine if HNQ can stimulate EET in *E. coli*, we tested an MG1655-derived BioDesignER (BioD)^40^ strain’s ability to generate current in a bioelectrochemical system (BES) using glycerol as an electron donor and HNQ as a redox shuttle. Under resting conditions, adding glycerol to the cells with HNQ resulted in immediate current generation (Figure S1A). In the presence of HNQ, BioD consumed 1.33 ± 0.03 mM glycerol and showed a peak current level of 101.8 ± 4.7 mA/m^2^. In contrast, without HNQ, BioD consumed 1.51 ± 0.04 mM glycerol and registered a peak current level of only 0.68 ± 0.02 mA/m^2^ (Figures S1A and 1B)^41^. To validate the redox cycling of HNQ, we performed a cyclic voltammogram analysis on BioD with HNQ and DMSO. The cyclic voltammetry of BioD showed that HNQ behaves like a redox shuttle and has a midpoint potential of -323 mV (Figure S1C). Consistent with prior studies, HNQ as a redox shuttle enables *E. coli* to perform EET using an anode as an extracellular electron sink.

To validate that the *E. coli* strains depend on oxidizing electron donors to generate EET using HNQ (Figure 1A), we deleted the three glycerol consumption routes (Figure 1B and S1D)^42^ to create a GlyNull strain (Figure S1E). We then compared the ability of BioD and GlyNull to generate EET and oxidize glycerol in a BES under resting conditions. BioD consumed 1.30 ± 0.04 mM glycerol and showed steady-state current levels of 74.8 ± 2.1 mA/m^2^. In contrast, GlyNull consumed ∼ one-tenth as much glycerol (0.141 ± 0.033 mM) and recorded only 2.3 ± 0.72 mA/m^2^ (Figures 1C and 1D). These results strongly suggest that GlyNull does not perform EET due to a lack of glycerol consumption. To confirm this loss of function was not due to polar effects, we reintroduced the *gldA* gene at its native genomic locus under a constitutive synthetic promoter (Figure S1E, Methods). This GldA+ strain exhibited restored glycerol consumption and EET levels, consuming 0.85 ± 0.07 mM glycerol and generating a steady-state current density of 54 ± 2 mA/m^2^ (Figures 1C and 1D). This data indicates that oxidation of a carbon source is crucial for HNQ-mediated EET.

Glycerol oxidation simultaneously initiates electron fluxes from the cytoplasm (NADH) and the inner membrane (reduced quinone). Thus, to simplify the investigation of the HNQ-mediated EET mechanism, we chose pyruvate as an electron donor rather than glycerol, as it would initiate electron fluxes only from the cytoplasm (Rationale in Methods). To validate the choice of pyruvate as the electron donor, we deleted the ubiquinone-dependent^43^ *poxB* gene to generate the PoxNull strain (Figure S1F) since PoxB oxidizes pyruvate and reduces the quinone pool. We then investigated PoxNull’s ability to generate EET with complete consumption of the provided pyruvate and glycerol as the control electron donor. The PoxNull strain consumed nearly all the provided pyruvate while consuming only 25% of the glycerol (Figure S1H). Although glycerol is a more reduced electron donor than pyruvate and is expected to stimulate higher EET, the PoxNull strain generated similar EET levels with glycerol or pyruvate as electron donors (Figure S1G). This result indicates that pyruvate is a more suitable choice of electron donor for investigating the EET mechanism.

### *E. coli* upregulates diverse redox pathways in response to HNQ

With data supporting that HNQ-dependent EET is linked to catabolism, we next sought to identify the biomolecule that catabolism reduces and serves as an electron donor to HNQ. Such identification is often recognized as a formidable challenge because catabolic pathways are intricately linked to many redox biomolecules. To address this challenge, we first profiled the transcriptome of HNQ-treated cells under no EET (BES) condition to infer which cellular redox pools HNQ engages. The HNQ-treated cells show only 2.1% differentially expressed genes (DEG), yet perturbed genes involve several redox biomolecules (Figure 1E). Upregulated gene transcripts include pathways for putative naphthoquinone degradation, naphthoquinone sensing transcription factors, and enzymes that oxidize NAD(P)H (Figure 1F). Regulon-level transcriptional analysis suggested simulation of the oxidative stress regulator OxyR^44,45^ (Figures 1G and S1I). Within central metabolism, HNQ upregulates the expression of enzyme subunits involved in pyruvate dehydrogenase and the oxidative TCA cycle (Figure S1J). The transcriptomics results encouraged us to investigate all the cellular electron carriers under EET conditions to elucidate the mechanism of HNQ-mediated EET.

### NADH is a key enabler of HNQ-mediated EET

As the redox perturbations induced by HNQ were broad, we set out to experimentally probe each of the electron carriers (NADH, NADPH, inner membrane quinones, glutathione, and thioredoxin) that could possibly interact with the HNQ-mediated EET pathway (Figure 1A). Since these electron carriers are interconnected, perturbing any of these electron carriers could alter the EET levels. However, we aimed to pinpoint the exact redox biomolecule that reduces HNQ through extensive deletions of redox-related genes. Thus, we established the following criteria to assess the outcome of perturbing the redox pool levels: (1) Gain in EET levels suggests that the deleted gene(s) compete with HNQ for electron fluxes, (2) Less than 50% loss in EET levels suggests that the deleted gene(s) do not directly reduce HNQ and is instead a pleiotropic effect, (3) More than 50% loss in EET levels indicates that the deleted gene(s) directly reduce HNQ. The fold change in the total charge deposited on the anode by the mutant and parent strains was utilized to assess the extent of EET gain or loss.

Our first line of inquiry explored the possibility of the inner membrane respiratory reductases reducing HNQ (Figure 2A). The anaerobic respiratory reductases can reduce naphthoquinones *in vitro*^46,47^. To understand if these reductases can reduce HNQ *in vivo*, we generated the AnoxicNull strain lacking native anaerobic respiratory pathways (Figure 2B, top). To confirm the phenotype, we show that AnoxicNull does not benefit anaerobic growth while respiring on the electron acceptors (Figure S2A). Next, we evaluated AnoxicNull’s ability to produce current with HNQ compared to BioD under resting conditions. AnoxicNull showed a gain in current density peak with 90.6 ± 3.8 mA/m^2^, whereas BioD generated a current density peak of 67.7 ± 5.4 mA/m^2^ (Figure 2B, bottom). AnoxicNull deposited a 1.7-fold higher charge on the anode than BioD (Figure S2B). These data support that respiratory reductases do not reduce HNQ and instead suggest that the respiratory reductases compete with HNQ for electrons.

**Figure 2:**
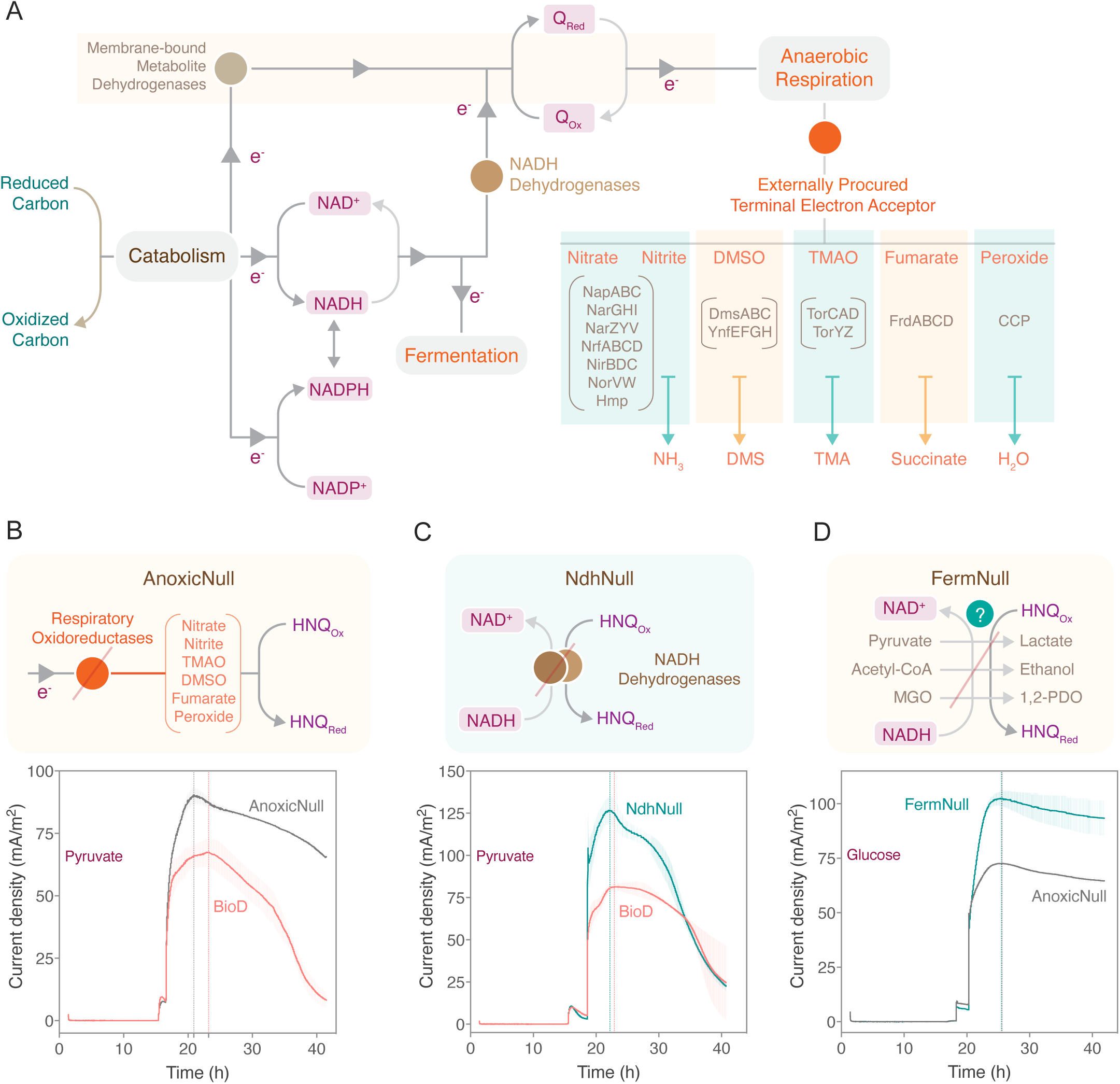
NADH is critical for HNQ-mediated EET. (A) A schematic illustrating electron flow from the carbon catabolism into the anaerobic respiratory and fermentative pathways. Diverse cellular electron carriers like the NADH pool, the NADPH pool, and the quinone pool orchestrate this electron flow into different electron sinks. (B) (top) The *E. coli* respiratory oxidoreductases that reduce the known terminal electron acceptors are deleted in the AnoxicNull strain. (bottom) Chronoamperometry of the AnoxicNull and BioD strains under resting conditions, showing that the peak and steady-state current density in AnoxicNull is higher than that in BioD. This data suggests that enzymes responsible for nitrate, nitrite, nitric oxide, DMSO, TMAO, fumarate, and peroxide reduction do not distinctly reduce HNQ. The data represent mean ± standard deviation obtained in triplicate and duplicate BES experiments of BioD and AnoxicNull, respectively. (C) (top) The *E. coli* NADH dehydrogenases that transfer electrons from NADH to quinones are deleted in the NdhNull strain. (bottom) Under resting conditions, NdhNull has a higher peak current density than the BioD strain, suggesting that the NADH dehydrogenases do not reduce HNQ. (D) (top) The *E. coli* fermentative enzymes that transfer electrons from NADH to pyruvate, acetyl-CoA, and MGO are deleted in the FermNull strain. (bottom) Under resting conditions, FermNull maintains higher peak and steady current density levels than the AnoxicNull strain. This data indicates that higher availability of NADH enhances the HNQ-mediated EET. The strains in this experiment were supplemented with 20 mM glucose in the biomass-generating culture and tested with 2 mM glucose as the electron donor in the BES. 20 mM pyruvate and 20 mM glycerol were added to the biomass-generating culture of the strains investigated in (B) and (C). 2 mM of pyruvate as the electron donor was added in the BES to stimulate EET. The data in (C) and (D) represent mean ± standard deviation obtained in triplicate BES experiments.

Naphthoquinones can be reduced by non-respiratory oxidoreductases in the inner membrane that partner with menaquinone and by the menaquinone pool in a non-enzymatic fashion (Figures 2A and S2C)^48^. To understand if the menaquinone pool significantly assists in reducing HNQ, we generated the MqNull strain by deleting the menaquinone biosynthetic pathway (Figure S2D). We then tested MqNull’s ability to produce current using HNQ alongside BioD under resting conditions. MqNull strain generated a higher peak current density of 91.3 ± 5.5 mA/m^2^ than the BioD strain (74.6 ± 0.9 mA/m^2^), although it showed only a minor difference in the overall charge deposited on the anode (Figure S2E). This data indicates that the menaquinone pool may weakly compete for electron fluxes with the HNQ-mediated EET pathway.

We next sought to investigate if NADH dehydrogenases directly reduce HNQ (Figure 2A). NADH dehydrogenase reduces redox shuttles in other mediated EETs^38,49,50^. Moreover, naphthoquinones like HNQ are structurally similar to menaquinone and are known to be reduced by NADH dehydrogenases^51–54^. To elucidate if NADH dehydrogenases enable HNQ-mediated EET in *E. coli*, we generated the NdhNull strain that does not encode the two NADH dehydrogenases NDH-I and NDH-II in *E. coli* (Figures 2C, top, and S2F-G). We then tested the HNQ-mediated EET in NdhNull and BioD under resting conditions. The NdhNull strain generated a 1.3-fold higher charge deposition on the anode than the BioD strain (Figure S2H). The NdhNull showed a higher current density peak (127 ± 7.8 mA/m^2^) than BioD (81.4 ± 1.8 mA/m^2^) (Figure 2C, bottom). These data indicate that NADH dehydrogenases NDH-I and NDH-II do not reduce HNQ. Importantly, AnoxicNull, MqNull, and NdhNull, designed to have limited electron fluxes from the NADH pool into the inner membrane, show higher EET output, strongly suggesting that the electron donor(s) to HNQ are present in the cell cytoplasm.

To further validate NADH’s involvement, we hypothesized that deleting competing NADH sinks within fermentative pathways would boost the electron flux through the HNQ-mediated EET (Figure 2A)^55,56^. To test this hypothesis, we deleted the enzymes catalyzing the formation of lactate, ethanol, and 1,2-propanediol in AnoxicNull to generate the FermNull strain (Figures 2D, top, and S2I). To test the EET performances of FermNull and AnoxicNull, we chose glucose over pyruvate as the electron donor since it yields more NADH. Under resting conditions, FermNull generated a 1.4-fold higher charge deposition on the anode than AnoxicNull (Figure S2J). FermNull showed a higher current density peak (102.6 ± 3.4 mA/m^2^) than AnoxicNull (72.6 ± 0.6 mA/m^2^) (Figures 2D, bottom). Interestingly, both these strains showed similar formate and acetate profiles at the end of the BES run (Figure S2K). Together with prior data (Figure 2C), these results strongly suggest that NADH supplies electrons to the HNQ-mediated EET.

### NADPH, glutathione, and thioredoxin are not the key players in HNQ-mediated EET

Having established that NADH is important for reducing HNQ, we next examined the relationship of NADPH with HNQ-mediated EET. While NADPH is mandatory for cell growth (Figure 3A, left), cells also use its reductive power to detoxify electrophiles like naphthoquinones^57–59^. To probe if NADPH enables HNQ-mediated EET, we engineered AnoxicNull to generate the NADPHnull strain that can only generate NADPH when gluconate is present (Figure 3A, right, and 3B). To confirm the gluconate auxotrophy, we show that NADPHNull achieved aerobic growth with gluconate supplementation but not in the 2xYT media (Figure 3C). We then probed the EET performance of NADPHnull and AnoxicNull with pyruvate as the electron donor. NADPHnull with 52.8 ± 5.8 mA/m^2^ showed a lower peak current density than AnoxicNull (72.4 ± 0.6 mA/m^2^) (Figure 3D). However, the difference in the charge deposited on the anodes by NADPHnull (5.96 ± 0.64 coulombs) and AnoxicNull (6.14 ± 0.15 coulombs) was not significant (Figure S3A). This data strongly suggests that NADPH does not play a dominant role in reducing HNQ. We also investigated whether transhydrogenases can reduce HNQ via their promiscuous activity^60^. We deleted both the transhydrogenases in the AnoxicNull strain to generate a THnull strain (Figure S3B) and saw no notable loss of EET, indicating that SthA does not reduce HNQ (Figure S3C).

**Figure 3:**
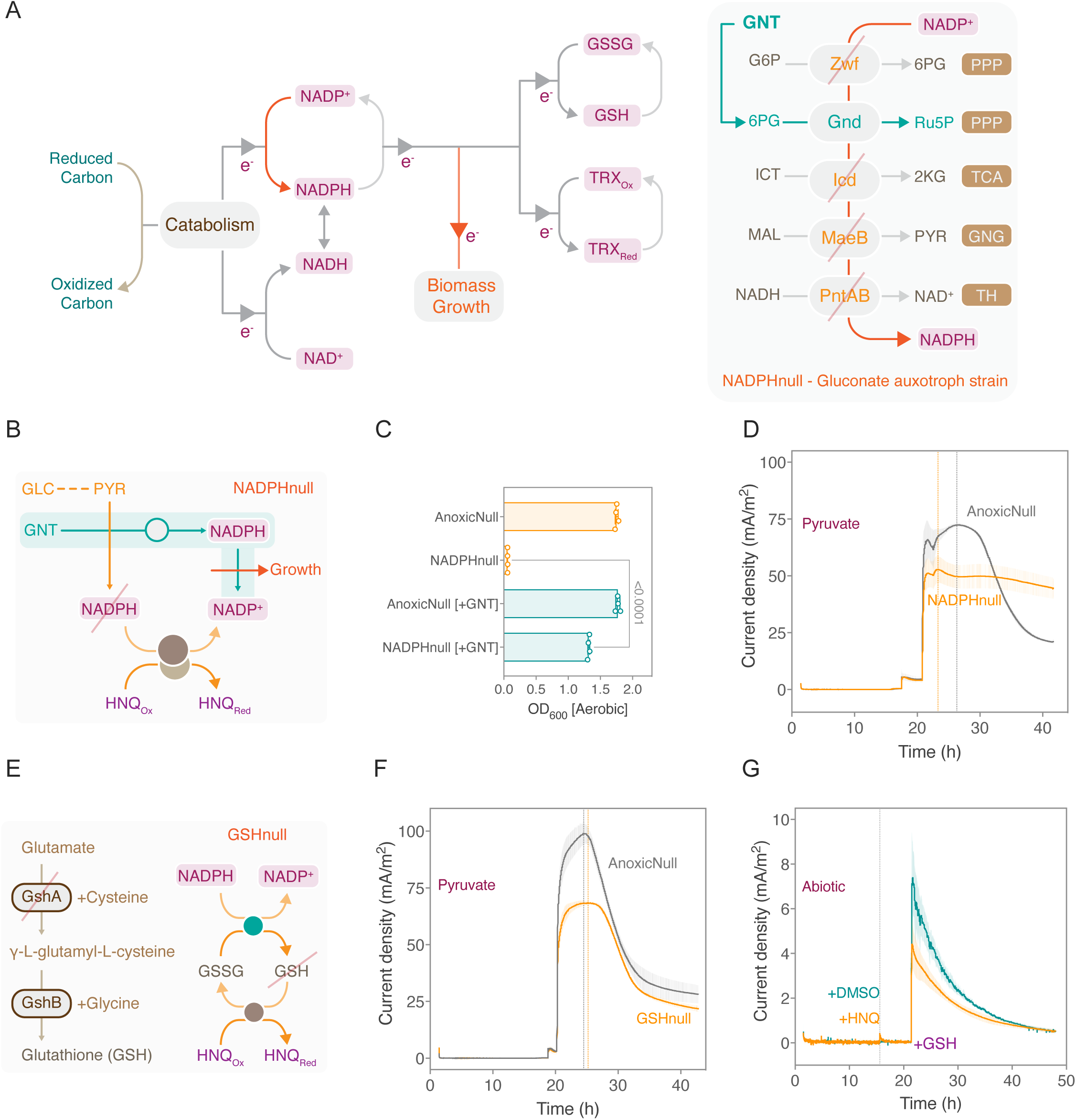
NADPH, glutathione, and thioredoxins are not critical for HNQ-mediated EET. (A) (Left) A schematic of cellular redox reactions powered by NADPH in *E. coli*. NADPH generated from carbon catabolism is indispensable for cell growth and global redox homeostasis. (Right) Eliminating NADPH-generating enzymes Zwf, Icd, PntAB, and MaeB renders the *E. coli*-NADPHnull strain into a gluconate (GNT) auxotroph. (Abbreviations: G6P, glucose 6-phosphate; 6PG, 6-phosphogluconate; Ru5P, ribulose 5-phosphate; ICT, D-threo-isocitrate; 2KG, α-ketoglutarate, MAL, malate; PYR, pyruvate; Zwf, Glucose-6-phosphate dehydrogenase; Gnd, 6-phosphogluconate dehydrogenase; Icd, Isocitrate dehydrogenase; MaeB, Malate dehydrogenase; PntAB, Pyridine nucleotide transhydrogenase, PPP, Pentose Phosphate Pathway; TCA, Tricarboxylic Acid cycle; GNG, Gluconeogenesis; TH, Transhydrogenase; GNT, Gluconate) *(B-D) The NADPH pool does not stimulate the HNQ-mediated EET pathway directly.* (B) A schematic to show that the NADPHnull strain is incapable of both NADPH production and growth when either pyruvate (PYR) or glucose (GLC) is the carbon source. However, cells can generate NADPH and grow biomass with gluconate (GNT) as the carbon source. (C) OD_600_ of AnoxicNull and NADPHnull strains after 16 hours of aerobic growth, showing that NADPHnull does not grow on 2x-YT media, whereas NADPHnull strain accumulated cell biomass in 20 mM gluconate-supplemented 2x-YT media. This data confirms that NADPHnull is a gluconate auxotroph. The strains in the experiment were pre-grown in LB supplemented with 20 mM of gluconate and 1 mM MgSO_4_. The data represent mean ± standard deviation obtained in four biological replicates. The P-value is calculated using the Welch’s t-test with a significance threshold of 0.05. (D) Chronoamperometry of NADPHnull and AnoxicNull strains under resting conditions, showing NADPHnull reached saturation current levels earlier than AnoxicNull, although with a smaller value of peak current density. The strains in this experiment were supplemented with 20 mM gluconate and 20 mM pyruvate in the biomass-generating culture. The data represent mean ± standard deviation obtained in triplicate BES experiments. *(E-G) Eliminating glutathione biosynthesis has a pleiotropic effect on HNQ-mediated EET* (E) (Left) An illustration of the glutathione biosynthetic pathway in *E. coli*. Deletion of *gshA* abolishes GSH biosynthesis. (Right) GSH is known to reduce HNQ both enzymatically and non-enzymatically. (F) Under resting conditions, GSHnull generates a lower current density than the AnoxicNull strain. The attenuated current level is attributed to a trickle-down effect of the *gshA* deletion. The data represent mean ± standard deviation obtained in triplicate BES experiments. (G) Chronoamperometry investigation of abiotic electron transfer from GSH to HNQ, showing that 40 µM GSH is oxidized at the anode poised at 200 mV vs. Ag/AgCl with and without 20 µM HNQ. This data suggests that GSH does not influence HNQ-mediated EET via abiotic electron transfer. The data represent mean ± standard deviation obtained in triplicate BES experiments. The strains in (F) and (G) were supplemented with 20 mM pyruvate and 20 mM glycerol in the biomass-generating culture. The EET levels in (F) were tested in BES with 2 mM of pyruvate as the electron donor.

The glutathione^61–66^ and the protein thiols^67–70^ are known to reduce naphthoquinones enzymatically and form adducts^71^. We also found that HNQ stimulates the upregulation of protein thiols like *aphC* and *trxC* (Figure 1F). Thus, we hypothesize that glutathione and protein thiols like thioredoxins can reduce HNQ and enable EET (Figures 3A and 3E). To elucidate the role of GSH in HNQ-mediated EET, we generated the GSHnull strain by deleting the first gene *gshA* of the GSH biosynthetic pathway (Figure S3D). We then compared the EET performance of the AnoxicNull and the GSHnull strains under resting conditions. The GSHnull strain deposited a 0.8-fold lower charge on the anode than the AnoxicNull strain (Figure S3E). GSHnull showed a smaller current density peak (68.4 ± 0.7 mA/m^2^) than AnoxicNull (99 ± 4.8 mA/m^2^) (Figure 3F). With our defined rubrics, this data suggests that the GSH has a pleiotropic effect on HNQ-mediated EET. However, there is also a possibility that GSH abiotically reduces HNQ. To test if GSH reduces HNQ abiotically (Figure S3F), we mixed GSH and HNQ in a BES and monitored the current. In this abiotic chronoamperometry experiment, adding 40 µM GSH to HNQ and DMSO gave a peak current density of 4.4 ± 1.1 mA/m^2^ and 7.4 ± 1.5 mA/m^2^, respectively (Figure 3G). This data indicates that GSH is directly oxidized by the anode poised at 200 mV, regardless of the presence of HNQ. Although this result does not efficiently probe the abiotic reduction of HNQ by GSH, we conclude that the partial loss of EET in GSHnull is due to a pleiotropic effect.

Further, to elucidate the role of TRX in HNQ-mediated EET, we generated the TRXnull strain by deleting *trxA, trxB,* and *trxC* genes in the AnoxicNull strain (Figures 3A and S3G-H). We then compared the EET performance of the TRXnull and the AnoxinNull strains under resting conditions. We saw no loss of EET in the TRXnull strain, indicating that the thioredoxin pool does not reduce HNQ (Figure S3I).

### Cytoplasmic quinone reductases facilitate HNQ-mediated EET under resting conditions

The experimental observations so far suggest that NADH assists in reducing HNQ, and other cellular electron carriers like inner membrane quinones, NADPH, glutathione, and thioredoxin do not directly reduce HNQ. Thus, we hypothesized that NADH-utilizing quinone reductases may catalyze HNQ-mediated EET. *E. coli* hosts several such quinone reductases (QReds)^72–87^, and these QReds are essential players in xenobiotic metabolism^88^. To understand the role of quinone reductases in reducing HNQ, we generated the QRedNull strain by deleting several characterized and putative quinone reductases in the AnoxicNull strain (Figure 4A). We then compared the EET performance of QRedNull and AnoxicNull with different electron donors under resting conditions. QRedNull showed a current density peak of 13.2 ± 1.9 mA/m^2^, about 7.5-fold lower than AnoxicNull’s 98.3 ± 1.7 mA/m^2^, using 2 mM pyruvate as the electron donor. With an additional 1 mM glucose, QRedNull continued to maintain the current density level of 13.8 ± 0.3 mA/m^2^, whereas AnoxicNull generated 97.6 ± 2.9 mA/m^2^, indicating the substantial loss of EET extends to other electron donors (Figure 4B). This result indicates that QReds reduce HNQ.

**Figure 4:**
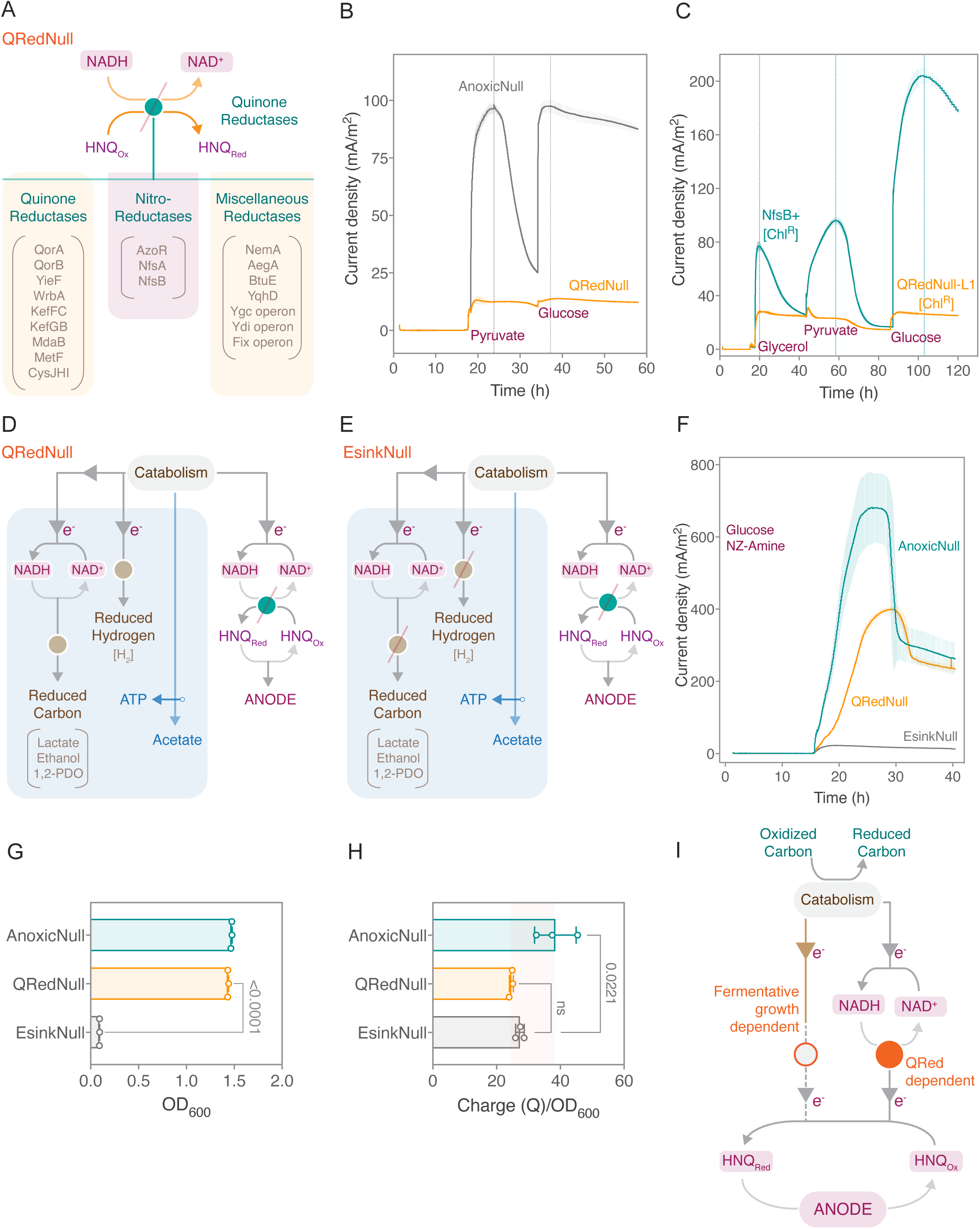
Nitroreductase NfsB enables HNQ-mediated EET under resting condition. (A) A schematic showing that the QRedNull strain lacks the known and the putative quinone reductases in *E. coli*. (B) Chronoamperometry of QRedNull and AnoxicNull strains under resting conditions. QRedNull produces insignificant current density levels compared to AnoxicNull, suggesting that quinone reductases facilitate HNQ-mediated EET. The strains in this experiment were supplemented with 20 mM glycerol and 20 mM pyruvate in the biomass-generating culture. The cells were first provided with 2 mM pyruvate as the electron donor in the BES. Once the pyruvate-powered current levels dropped for the AnoxicNull strain, 1 mM glucose was injected into the BES. The data represent mean ± standard deviation obtained in duplicate BES experiments. (C) Chronoamperometry of the NfsB^+^ and QRedNull-L1 strains under resting conditions, demonstrating that plasmid complementation of NfsB restores the HNQ-mediated EET in the QRedNull-L1 strain. The HNQ-mediated EET restoration is seen with 2 M each of glycerol, pyruvate, and glucose as carbon sources. The data represent mean ± standard deviation obtained in triplicate BES experiments. (D) The QRedNull strain can grow on fermentative pathways and does not perform quinone reductase facilitated HNQ-mediated EET. (E) The EsinkNull strain cannot grow on fermentative pathways and does not perform quinone reductase facilitated HNQ-mediated EET. (F) The EsinkNull strain generates insignificant current density levels compared to the AnoxicNull and the QRedNull strains. The loss of EET in EsinkNull is due to blocked electron flow into both the HNQ-mediated EET and the native fermentative pathways. The data represent mean ± standard deviation obtained in triplicate BES experiments. (G) OD_600_ of the AnoxicNull, the QRedNull, and the EsinkNull strains under growth conditions, showing that the EsinkNull strain cannot grow in anaerobic conditions. The data represent mean ± standard deviation obtained in triplicate BES experiments. The P-value is calculated using the Welch’s t-test with a significance threshold of 0.05.(H) The Charge (Q) to OD_600_ ratio of the QRedNull and the EsinkNull strain, showing equivalent charge transfer onto anode per unit cell of both these strains. The data represent mean ± standard deviation obtained in triplicate BES experiments. The P-values are calculated using one-way ANOVA, with a significance threshold of 0.05. (I) Proposed working model of HNQ-mediated EET in *E. coli*.

Knowing that QReds are central to HNQ-mediated EET, we investigated which specific QReds reduce HNQ. The DEG analysis of HNQ-treated cells previously showed upregulated *azoR* expression levels (Figures 1E-F). Thus, we first deleted *azoR* in AnoxicNull to generate the AzoNull strain. AzoNull showed only a minor loss of EET compared to AnoxicNull (Figure S4A)^89^. Next, we deleted the nitroreductases *nfsA* and *nfsB* in the AzoNull strain^90^, leading to a significant loss of EET compared to AnoxicNull (Figure S4A). Literature reports have established that NfsA activity is NADPH-dependent^91^, whereas NfsB is compatible with both NADH and NADPH as electron donors^92^. Thus, based on the conditions under which we performed the experiments, we concluded that NfsB is likely the key QRed facilitating HNQ-mediated EET.

Next, to confirm the role of NfsB, we reintroduced the *nfsB* gene under the P_ara.AM_ promoter on a low copy plasmid (Figure S4B). The NfsB^+^ strain generated peak current densities of 77.1 ± 3.0 mA/m^2^ with 2 mM glycerol, 96.0 ± 2.4 mA/m^2^ with 2 mM pyruvate, and 204.3 ± 5.5 mA/m^2^ with 2 mM glucose, restoring the EET levels in the QRedNull-L1 strain (Figure 4C). The NfsB+ strain deposited 1.8, 2.7, and 6.6 fold higher charge than QRedNull-L1 with glycerol, pyruvate, and glucose as the electron donors, respectively (Figure S4C). This data illustrates that the nitroreductase NfsB facilitates the HNQ-mediated EET by oxidizing NADH. In summary, HNQ-mediated EET occurs through cytoplasmic HNQ-reductases under resting conditions. Moreover, HNQ-mediated EET restores NAD^+^ regeneration with the anode as an electron sink, the hallmark of a respiratory pathway.

### Additional routes of HNQ-mediated EET exist under fermentative growth conditions

Having established the HNQ-mediated EET mechanism under resting conditions, we aimed to elucidate the behavior of *E. coli* HNQ-mediated EET during growth conditions. We postulated that, under anaerobic growth conditions, additional NAD^+^ regenerating HNQ-reductases might be expressed. To identify any additional HNQ-reductases, we eliminated fermentative pathways that serve as confounding factors by facilitating NAD^+^ regeneration during anaerobic growth. Consequently, we generated the EsinkNull strain by deleting the fermentative pathways in QRedNull (Figures 4D-E). We then tested the EET output of the AnoxicNull, the QRedNull, and the EsinkNull strains under growth conditions (Rationale in Methods). The AnoxicNull, the QRedNull, and the EsinkNull strains showed current density peaks of 682 ± 9.1 mA/m^2^, 400.6 ± 9.3 mA/m^2^, and 22 ± 1.4 mA/m^2^ respectively (Figure 4F). This data strongly indicates that EsinkNull no longer contains a pathway for HNQ-mediated EET. AnoxicNull and QRedNull oxidized all the provided glucose to achieve anaerobic growth and perform EET (Figures 4G and S4F-I). Even though EsinkNull is metabolically active (Figure S4I), it oxidized ∼10-fold less glucose than the other strains, showed no drop in the pH, and did not show biomass growth (Figures 4G and S4E-F). QRedNull, on the other hand, showed 0.6-fold and 14.1-fold charge deposition on the anode compared to AnoxicNull and EsinkNull, respectively (Figure S4D). These observations suggest that the fermentative growth in QRedNull also enables cells to perform HNQ-mediated EET. Intriguingly, the difference in the values of charge deposited on the anode per unit cell of the QRedNull and the EsinkNull strains was insignificant (Figure 4H). This analysis indicates that a baseline HNQ-mediated EET occurs per unit cell independent of QReds under growth conditions.

To address the source of the EET output of QRedNull under growth conditions, we hypothesized that the enzymes involved in formate and hydrogen metabolism could reduce HNQ. Hence, we deleted the three formate dehydrogenase operons in BioD to generate the FDHnull strain and tested its EET performance alongside BioD under resting conditions^93^. BioD and FDHnull showed similar peak current density levels with pyruvate as the electron donor. However, BioD maintained a 6-fold higher steady-state current density than FDHnull (Figures S4J-K). Additionally, metabolite measurements of spent media showed that BioD had ∼0.6-fold less formate than FDHnull (Figure S4L). These results suggest that some EET output comes from the formate/hydrogen metabolism, which proportionally increases during the BES run with cell growth, as we see in QRedNull. Based on all the experimental evidence, we propose that selective electron transfer from the NADH pool to HNQ via quinone reductases facilitates HNQ-mediated anode respiration. At the same time, additional route(s) of EET is coupled with fermentative growth (Figure 4I).

### EET in growing cells is characterized by redox and stress expression perturbations

We next sought to determine the gene expression changes associated with EET in both growing and non-growing strains. We hypothesized that the gene expression changes between the constructed strains would further demonstrate the interconnectivity of anaerobic electron sink, redox stress, and biomass growth. Unfortunately, harvesting cells from BES is logistically challenging since cells localize in the anode during the chronoamperometry run. Thus, to efficiently collect cells to perform transcriptomics, we designed a representative colorimetric assay performed in the anaerobic chamber that investigates HNQ-mediated EET using amaranth, a redox dye, as an electron sink (Rationale and details in Methods).

Using the amaranth assay, we profiled the gene expression of AnoxicNull (which has both fermentative and NfsB-based routes to reduce HNQ), QRedNull (which has only fermentation), and EsinkNull (which has neither) in the presence of HNQ under growth conditions (Figure 5A and S5A-C). A clear separation between gene expression across the strains was observed, with EsinkNull being farther separated from both AnoxicNull and QRedNull, consistent with the fact that EsinkNull is unable to grow (Figure 5B). Transcriptional differences across strains are driven by a down-regulation of growth pathways (ppGpp, translation, thiamine, pyrimidine, arginine regulons) and up-regulation of stress (rpoS, soxS regulon) and energy starvation (Crp-1) pathways in the non-growing EsinkNull strain, consistent with a well-characterized growth/stress transcriptional tradeoff in *E. coli*, termed the ‘fear/greed’ tradeoff (Figure 5B)^94^.

**Figure 5:**
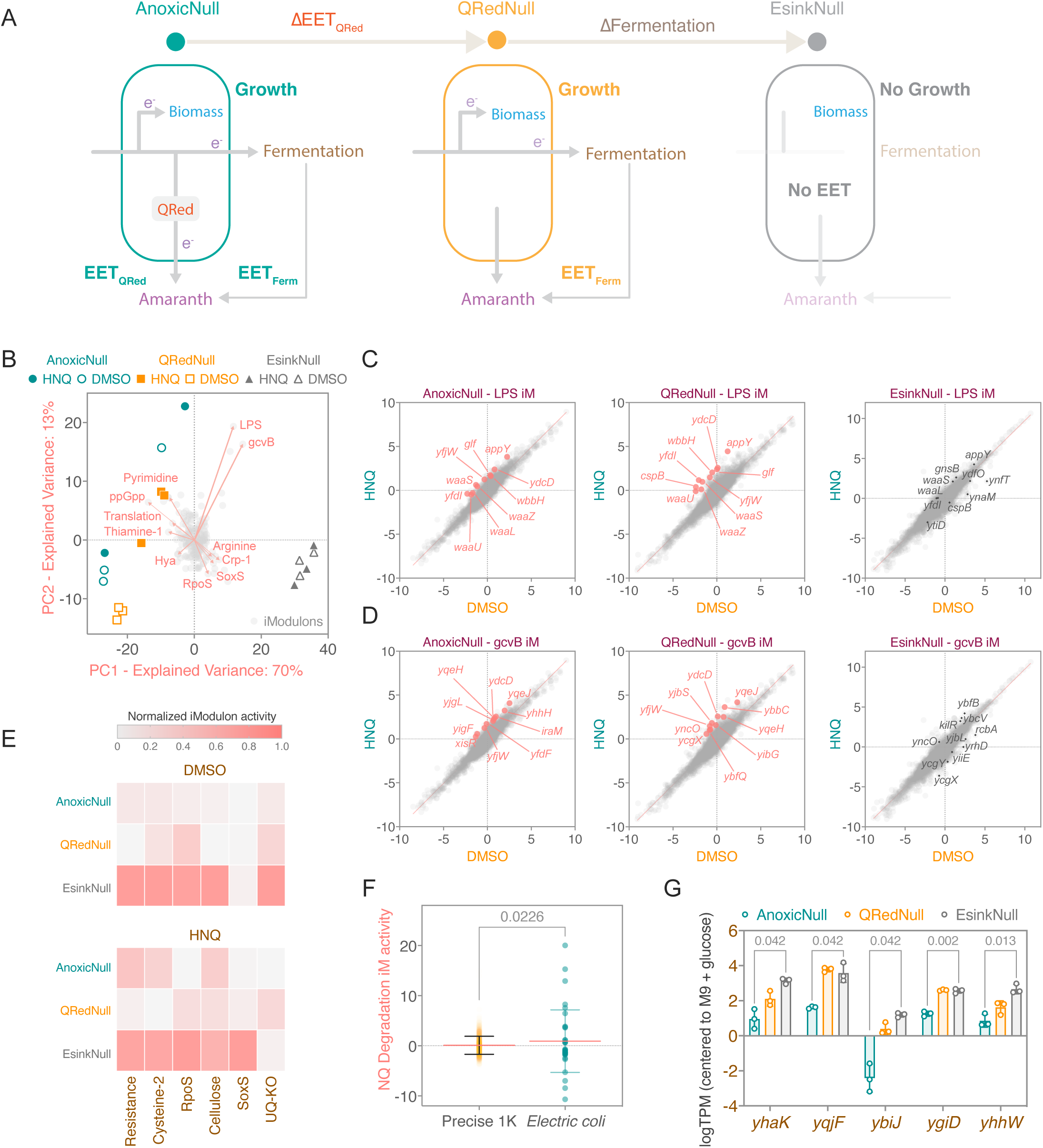
Transcriptomics analysis reveals HNQ-induced response under growing conditions. (A) The strains with varied growth and EET capabilities are tested on their ability to reduce amaranth dye using HNQ-mediated EET under anaerobic conditions (Details in Methods). (B) Biplot for principal component analysis of the activity matrix **A** computed with Independent Component (iModulon) Analysis for AnoxicNull, QRedNull, and ESinkNull with and without HNQ supplementation. (C) Different gene expression plots for AnoxicNull, QRedNull, and ESinkNull, with highly perturbed genes within the LPS iModulon highlighted. Gene expression was centered on an aerobic glucose reference condition. (D) Different gene expression plots for AnoxicNull, QRedNull, and ESinkNull, with highly perturbed genes within the gcvB iModulon highlighted. Gene expression was centered on an aerobic glucose reference condition. (E) Heatmap of redox stress iModulon activities across strains and conditions. (F) Boxplot of expression of a naphthoquinone degradation iModulon across an *E. coli* expression database (PRECISE1K) and the data generated within this project (*Electric coli*). The broad perturbation of genes within this iModulon in the project-specific data (“*Electric coli*”) resulted in this new iModulon appearing when ICA was run on PRECISE1k with the project data added. (G) Expression of genes within the naphthoquinone degradation iModulon. All the P-values are calculated using Welch’s t-test, with a significance threshold of 0.05.

The introduction of HNQ had a pronounced and similar effect on AnoxicNull and QRedNull. In these strains, HNQ supplementation reduced rpoS activity and perturbed two primary transcription modules (Figure 5B). The first perturbation was a down-regulation of the hya operon, which is associated with redox response and microaerobic growth conditions^45^. The second perturbation was a pronounced activation of a set of poorly characterized transcriptional modules containing lipopolysaccharide (LPS) genes (Figure 5C), gcvB-associated genes (Figures 5D), membrane-associated genes, prophage genes, and a large number of y-genes (Figures S5G-H and S5L). While the functional association to HNQ is unclear, these LPS and gcvB-associated transcriptional modules show a high degree of correlation across an *E. coli* gene expression data compendium and are activated under redox stress-related conditions^45^ (Figure S5K). Notably, the gcvB regulon was downregulated on pyruvate under non-growing conditions with HNQ, and LPS was not significantly changed, indicating that this activation is specific to growing conditions in the presence of HNQ (Figure 1G).

While AnoxicNull and QRedNull generally exhibited similar responses to HNQ, there were certain additional strain-specific responses to HNQ. In the AnoxicNull strain, the NAD biosynthesis gene nadB was downregulated in response to HNQ, suggesting an interaction between HNQ and the NADH pool (Figure S5D). In the QRedNull strain, there was an HNQ-induced activation of a ubiquinone knockout-associated regulon that contains the deleted *nfsB* (Figures 5E, S5E and Sup. Table 2). In the ESinkNull strain, a set of genes associated with HNQ degradation is up-regulated potentially to mitigate HNQ-mediated stresses when EET cannot be carried out (Figures 5F-G). Additional activation of SoxS was observed that could indicate additional redox stress when cells are not able to perform EET (Figures S5F and S5I), and the acetate kinase-encoding gene *ackA* is upregulated (Figure S5J), consistent with measured acetate production under growth condition (Figure S4I).

### Model simulations predict that HNQ-mediated EET facilitates anaerobic growth

We then put forward the question of whether there is a rational basis for the use of HNQ reduction as a part of anaerobic growth. This question is well suited to be explored with genome-scale constraint-based metabolic modeling, which computes optimal metabolic states under defined conditions^95^. Flux balance analysis using the genome-scale metabolic model for *E. coli,* iML1515^96^, revealed distinct metabolic adaptations to genetic modifications across strains. Under anaerobic conditions with a glucose uptake rate of 10 mmol/gDW/hr, BioD and AnoxicNull demonstrated a wild-type-like growth rate of 0.15 hr^-1^, utilizing mixed acetate fermentation pathways. In stark contrast, with most NAD-regenerating enzymes removed and the transhydrogenase (pntAB) activity capped to WT levels, the FermNull and EsinkNull strains exhibited a severely reduced growth rate of 0.01 hr^-1^, utilizing the Entner-Doudoroff pathway to minimize NADH production and Malate Dehydrogenase for NAD^+^ regeneration. Strikingly, integrating a quinone reduction enzyme (QRed) into the FermNull strain restored growth to 0.19 hr^-1^, outperforming the WT strain (Figure 6A). Simulations indicate the growth advantage of HNQ-mediated anaerobic metabolism is largely due to the ability of the cell to secrete additional acetate rather than ethanol or lactate, which normally provide anaerobic routes to regenerate NAD, thereby enabling greater ATP production through acetate kinase (Figure S6A). Additionally, the inclusion of proteome allocation considerations through simulations with metabolism and macromolecular expression (ME) models suggested a reduction in both stress-related proteome and total proteome requirements in FermNull+nfsB compared to WT (Figures S6B-C)^97,98^. This analysis suggests that HNQ-mediated EET can not just rescue anaerobic growth in fermentation-impaired strains but may even offer a more efficient pathway both in terms of stoichiometric and proteomic efficiency, assuming the HNQ can be efficiently recycled.

**Figure 6:**
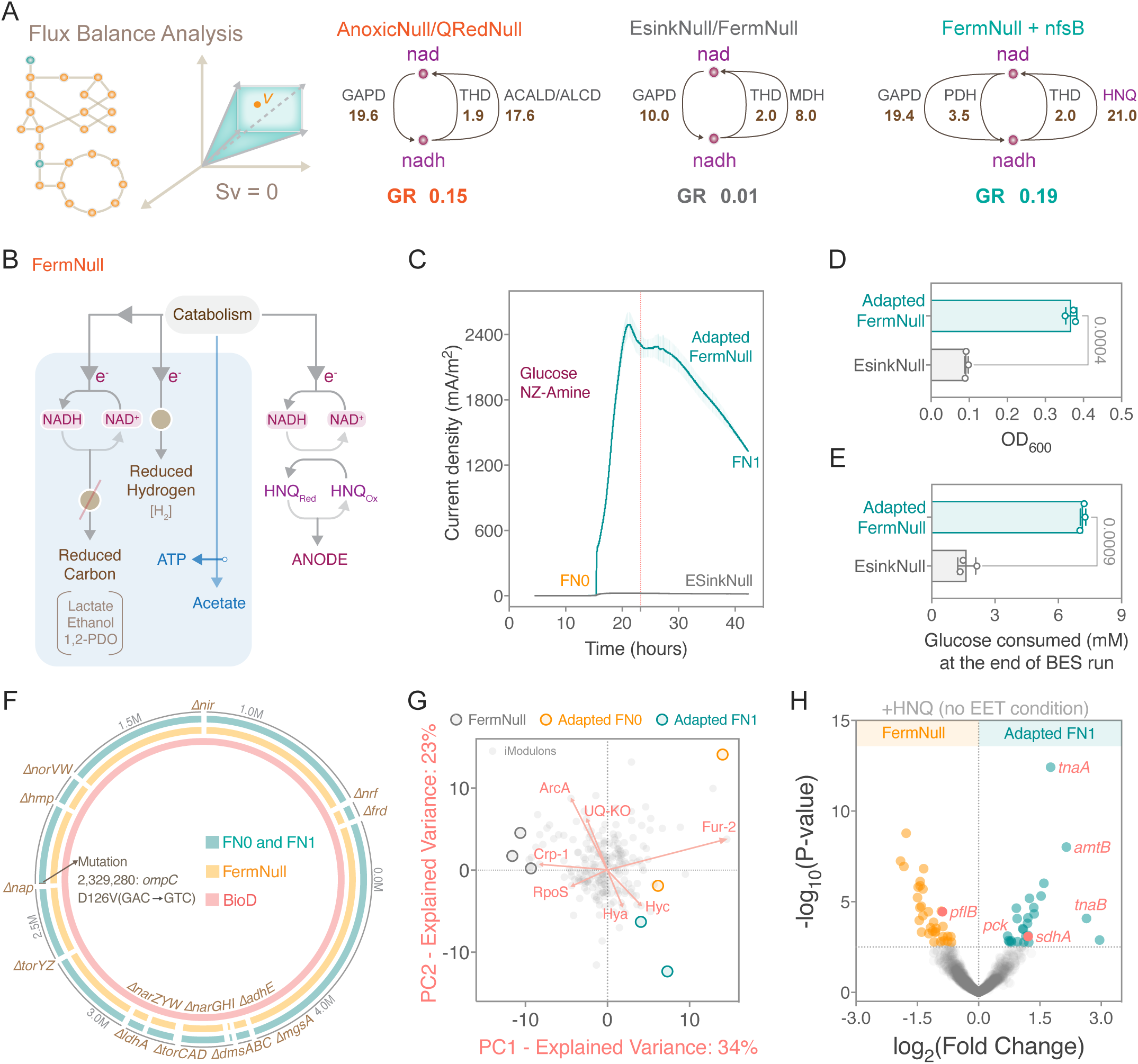
HNQ-mediated EET enables *E. coli* to respire and grow on anode. (A) Flux balance analysis of *E. coli* strains under anaerobic growth on glucose shows the recovery of anaerobic growth via naphthoquinone and NfsB. Fluxes in mmol/gDW/hr units, growth rate (GR) in 1/hr units. The listed fluxes represent >95% of the turnover of NAD/NADH. Reaction abbreviations - GAPD: Glyceraldehyde 3-phosphate Dehydrogenase. THD: Transhydrogenase. ACALD: Acetaldehyde Dehydrogenase. ALCD: Alcohol Dehydrogenase. MDH: Malate Dehydrogenase. PDH: Pyruvate Dehydrogenase. HNQ: Naphthoquinone Reductase (B) A schematic showing that the FermNull strain cannot grow on fermentative pathways; however, the quinone reductases are intact to enable HNQ-mediated anode respiration. (C) Chronoamperometry of the FN0 and the EsinkNull strains under growth conditions, showing that the FN0 strain carries over the EET phenotype gained in the prior extended BES run of the FermNull strain. (D) OD_600_ of FN0 and (E) Glucose oxidized by FN0 at the end of the BES run were significantly higher than EsinkNull. These results indicate that HNQ-mediated EET drives catabolism and biomass growth by respiring on the anode. The data in (C), (D), and (E) represent mean ± standard deviation obtained in triplicate BES experiments. The P-values are calculated using Welch’s t-test,, with a significance threshold of 0.05. (F) Genome diagram showing the sequences of BioD, FermNull, and FN0/FN1. Gene deletions in the FermNull strains are highlighted as gaps in the circular genome and the lone *ompC* mutation in FN0 and FN1 is highlighted. (G) Biplot for principal component analysis of the activity matrix **A** computed with Independent Component (iModulon) Analysis for the FermNull, FN0 and FN1 strains under no EET conditions. (H) Differential gene expression (DEG) analysis comparing FN1 to FermNull, showing perturbation of amino acid related genes *tnaAB* and *amtB* and anaplerosis and TCA-associated genes *pflB*, *pck*, and *sdhA*.

### *E. coli* adapts to BES conditions to grow on the anode as the sole respiratory electron sink

Based on modeling results, we postulated that HNQ-mediated EET could fuel an anaerobic energy metabolism without fermentation-driven NAD^+^ regeneration and restore growth in a fermentation-impaired *E. coli* strain. To elucidate if HNQ-mediated EET can conserve energy to enable growth in *E. coli*, we probed the FermNull strain that does not grow anaerobically but performs HNQ-mediated EET with QReds (Figure 6B). We tested the EET output of the FermNull and the EsinkNull strains under growth conditions. Surprisingly, the EET levels in FermNull significantly increased only after 39 hours (Figure S6D). FermNull showed a delayed peak current density of 1670 ± 124.9 mA/m^2^, whereas EsinkNull showed a consistent current density level lower than ∼23 mA/m^2^ (Figure S6D). The net charge deposited by FermNull by the end of the BES run was 55 folds higher than EsinkNull (Figure S6E). Thus, we labeled this strain as the adapted FermNull (FN0) strain. At the end of the BES run, FN0 showed a 6.6-fold higher OD_600_ than EsinkNull (Figure S6F). This data indicates that HNQ-mediated anode respiration facilitates biomass growth.

We passaged and saved a glycerol stock of the adapted-FermNull (FN0) strain from the BES run. We further probed the FN0 strain for EET and found that cells performed high levels of EET as soon as they were introduced in the BES. FN0 showed peak current density levels of 2493 ± 111 mA/m^2^, whereas EsinkNull showed a maximum current density of 24.2 ± 1.5 mA/m^2^ (Figure 6C). The FN0 strain showed about 4-fold higher growth compared to EsinkNull (Figure 6D). This high EET performance, coupled with growth, showed a drop in the pH of the BES spent medium (Figure S6G), consumed 4.3-fold higher glucose (Figure 6E), and generated only 0.12-fold acetate as compared to the EsinkNull strain (Figure S6H). We further passaged and saved a glycerol stock of FN0 at the end of the BES run and labeled it as FN1. A comparative analysis of DNAseq reads from FN1 and FN0 revealed a lone SNP mutation in the *ompC* gene when compared to the FermNull strain (Figure 6F). This is the first-ever demonstration of redox shuttle-mediated anaerobic growth on the anode as the sole respiratory electron sink.

We expression profiled FermNull, FN0, and FN1 to determine transcriptional changes associated with adaptation underlying improved growth under no EET condition. Two of the previously determined growth-associated fear transcription modules (RpoS, Crp) were higher in FermNull, consistent with previous results where these transcription modules were higher expressed on the non-growing ESinkNull strain (Figure 6G). The *hya* operon that we previously associated with HNQ supplementation and the *hyc* operon were both found to be higher in the adapted strains. In terms of new transcriptional modules, there were three modules most perturbed by the adaptation, specifically ArcA, Ubiquinone-KO, and Fur, indicating a possible derepression of the TCA cycle to generate more NADH. Notably, the HNQ-associated transcriptional module (LPS+gcvB) identified earlier was not perturbed by the growth adaptation. Looking at individual genes, *tnaAB*, *amtB*, *sdhA*, and *pckA* are up-regulated in the adapted strain, while *pflB* is down-regulated (Figure 6H). These perturbations to amino acid and anaplerotic metabolism genes indicate a metabolic reprogramming towards adapting to utilizing the provided peptide amino acids to sustain growth. Next, under amaranth assay conditions, FN0 showed 4.5-fold higher amaranth reduction than AnoxicNull (Figures S6I-K and S5A) and higher expression of gluconate incorporation enzymes and phosphoglycerate kinase as well as other amino acid perturbations (Figures S6L-M). FN0 also showed higher substrate-level phosphorylation activity in glycolysis (Figure S6N) than the increased acetate production predicted by the model-computed optimal metabolic state. Finally, we observed the FermNull strain exhibited strong activation of a NadR-associated NAD biosynthesis component (Figure S6O), and this perturbation was attenuated following adaptation (Figure S6P). These results suggest that *E. coli* can adapt to support EET-driven growth, driven by a general shift towards a greedy growth phenotype enabled by metabolic adaptation rather than upregulation of HNQ reductases.

## DISCUSSION

Here, we characterized a novel energy metabolism in *E. coli* in which redox cycling of HNQ enables respiration and growth on an extracellular anode. Through a reductionist approach, we found that a cytoplasmic quinone reductase, NfsB, enables HNQ-mediated EET by oxidizing NADH under resting conditions. Flux balance analysis predicted that HNQ-mediated EET can support anaerobic growth in *E. coli*, and this prediction was validated experimentally. Moreover, cells adapted to the BES conditions and generated A/m^2^ levels of EET output, two orders of magnitude higher than previously reported in *E. coli*. These adaptations were enabled by perturbed amino acid metabolism and activation of the anaplerosis-driven TCA cycle. Our findings reveal that *E. coli* has a latent extracellular respiratory pathway that drives anaerobic catabolism and growth.

In the following sections, we discuss the generalizable approach to examining the mechanisms of mediated EET pathways, along with the physiological significance and potential trade-offs associated with HNQ-mediated EET in *E. coli*.

### A blueprint for elucidating mechanisms and cellular objectives of mediated EETs

Unraveling the mechanisms of mediated EETs has been considered intractable^99,100^. Approximately one in six cellular enzymes in *E. coli* utilize cellular electron carriers. Thus, a large pool of candidate enzymes could reduce a redox shuttle like HNQ. Prior efforts to identify a partner have targeted certain redox-active enzymes using a combination of bottom-up genetics and *in vitro* biochemical methods^37,38,101–103^. However, these approaches have been collectively met with limited success. In this study, we introduce a radically different strategy leveraging the fact that the majority of the cellular electron flux passes through a limited number of cellular electron carriers: NADH, NADPH, ubiquinone, menaquinone, glutathione, and thioredoxins. We used genome editing to tactically alter electron fluxes through these carriers while maintaining cell viability under aerobic conditions. Subsequently, we examined these strains under anaerobic resting conditions to identify the key electron carrier enabling mediated EET. Testing EET under resting conditions was crucial for elucidating the electron carrier. Under resting conditions, we tracked the electron flux from a single electron source to electron sinks by measuring the EET output and the metabolite concentrations, allowing us to identify the key electron carrier. The specific electron carrier also alluded to the subcellular space where the redox shuttle is reduced, enabling us to narrow the search space for the enzymes capable of reducing the redox shuttle. From this narrowed list, we identified 1 of ∼300 possible redox enzymes responsible for reducing the majority of redox shuttle. We posit the methodology introduced here could be generalized to any combination of microbes and redox shuttles like phenazines, flavins, and quinones^100^.

This work also identified a minimal EET pathway that facilitates catabolism and growth. Although extracellular respiration on HNQ facilitates growth, not all redox shuttle and microbe combinations are expected to achieve growth-coupled EET. To identify such optimal combinations of microbes and redox shuttles, we propose leveraging a FermNull (Figure 6) equivalent strain of the microbe. As demonstrated in our study, we unraveled the emergent EET phenotype as an outcome of NADH oxidation by HNQ under BES conditions. This approach could be further augmented with the transcriptomic characterizations of cellular objectives under EET conditions, e.g., In the absence of an extracellular electron sink, we found HNQ stimulated the SoxS regulon. In summary, combining the above approaches will enable a holistic integration of mediated EETs with cell metabolism. It also establishes design principles for developing diverse electrical reporter modules for efficient biosensing and addressing redox imbalance challenges in bioelectrocatalysis. Additionally, these systems-level cell engineering strategies open new avenues for generating one-step reducing equivalents for electrosynthesis^104^ and minimal respiratory modules of synthetic minimal cells^105^.

### Physiological relevance of HNQ-mediated EET in *E. coli* and biosynthesis of HNQ

Cytoplasmic quinone reductases in association with HNQ could provide an economical mode of heterotrophy under nutrient-limited anaerobic conditions^106^. However, it raises a crucial question: what physiologically relevant terminal electron acceptors can HNQ reduce to benefit the anaerobic survival of microbes? Literature reports indicate reduced HNQ can abiotically transfer electrons to insoluble Fe(III)^107,108^, tellurite, selenite^109,110^, and azo dyes^111,112^. We speculate that *E. coli* could use soluble quinones like HNQ to respire on these diverse electron sinks in the gut and environmental niches. Similar naphthoquinone-mediated EET has been described in *Shewanella* species with humic acids as electron sinks^113–115^, as well as in lactic acid bacteria, *Klebsiella pneumoniae* L17, and even free-living fungi with extracellular Fe(III) as the electron sink^49,116–118^. Additionally, *E. coli* might utilize the endogenous HNQ as a respiratory electron sink, as previously shown with melanin in *Shewanella algae* BrY^119,120^.

Although HNQ is observed to reduce physiologically relevant extracellular electron sinks and is known to be secreted by *E. coli*^31,34^, we have yet to understand the physiological conditions necessary for HNQ biosynthesis in *E. coli*. Since HNQ shares structural similarities with menaquinone, it is likely derived from the DHNA biosynthetic pathway^121^. An understanding of HNQ biosynthesis in *E. coli* would give more critical insights into the physiological relevance of this novel energy metabolism. From an alternative perspective, *E. coli* can rely on obtaining HNQ from its ecological niche to perform an EET metabolism. Some exoelectrogens, like *Lactiplantibacillus plantarum,* utilize this communal resource strategy to perform EET without synthesizing the naphthoquinones they employ^122,123^. Although HNQ levels in ecological niches are unknown, it is a well-characterized secondary metabolite produced by various plants, facilitating interactions between plants and microbes^124,125^. In summary, *E. coli* could use HNQ-mediated energy metabolism in ecological niches with both endogenous and the communal presence of exogenous HNQ.

### Trade-offs of extracellular respiration mediated by HNQ

Given *E. coli’s* metabolic versatility, HNQ-mediated EET is expected to be selectively engaged under certain nutrient and environmental conditions due to its inherent trade-offs. Theoretically, as long as oxidized HNQ can be sufficiently regenerated, there are several motivating factors for *E. coli* to respire extracellularly, even when fermentation alone is possible. First, our constraint-based modeling suggests that EET is stoichiometrically and proteometrically more favorable than fermentation. Second, there is potentially less risk of growth-inhibiting fermentative metabolite accumulation. Third, the reliance on HNQ oxidation by external factors could create the basis for the community exchange of valuable metabolic resources, as predicted in other community models, leading to even more favorable metabolic scenarios^124,125^. Despite these benefits, HNQ-mediated EET entails two trade-offs. Firstly, HNQ does not oxidize the inner membrane quinone pool, eventually causing catabolic reactions involving inner membrane dehydrogenases to cease. Secondly, since HNQ-mediated EET does not utilize inner membrane reductases, it cannot exploit the proton-pumping activity of NADH dehydrogenase to generate additional proton motive force (PMF). Nevertheless, even with these trade-offs, we have demonstrated that HNQ enables extracellular respiration that drives catabolism and growth.

In conclusion, this study establishes a framework to unravel the molecular mechanism of redox shuttle-mediated EET. To further advance our understanding and engineering capabilities, future research must delve deeper into how redox shuttles interface with cellular redox biology to facilitate ATP and PMF generation. Key questions include whether the movement of reduced redox shuttles that do not require active transport across the inner membrane generates PMF. Additionally, the discovered HNQ-mediated EET paves the way for exploring other naphthoquinone-mediated extracellular respiration. It also lays a strong foundation for building a novel series of synthetic extracellular respiratory pathways that could modulate energy metabolism, gene expression, and overall cellular physiology.

## Acknowledgments

We thank Dr. Rob Egbert from the Pacific Northwest National Laboratory for providing the BioDesignER strain. We highly appreciate Dr. Steffen Lindner-Mehlich from the Max Planck Institute of Molecular Plant Physiology for valuable discussions on generating the NADPH auxotroph strain. We also thank Dr. Jayashree Soman from Rice University for proofreading the manuscript.

## Author contributions

Project Conceptualization: B.B.K, C.M.A-F;

Methodology: B.B.K, J.K, D.C.Z, C.M.A-F;

Investigation: B.B.K, R.S, J.K, A.P, D.C.Z;

Visualization: B.B.K;

Funding acquisition: C.M.A-F, B.O.P;

Supervision: C.M.A-F, D.C.Z, B.O.P;

Writing – original draft: B.B.K, J.K, R.S, A.P, D.C.Z, C.M.A-F;

Writing – review & editing: B.B.K, D.C.Z, C.M.A-F with inputs from everyone

---

**Supplementary Table 1:**
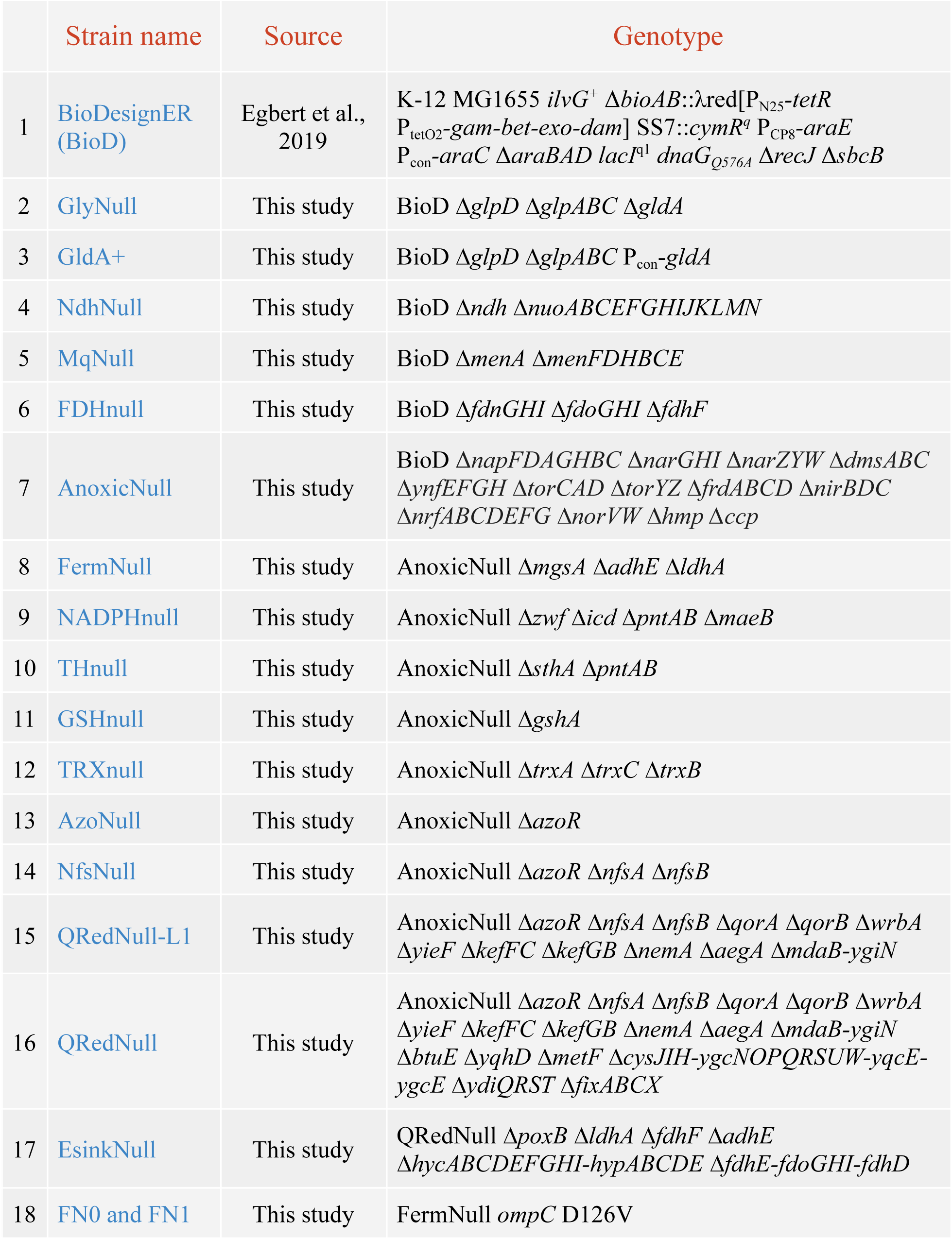
Strain List.

**Supplementary Table 2:**
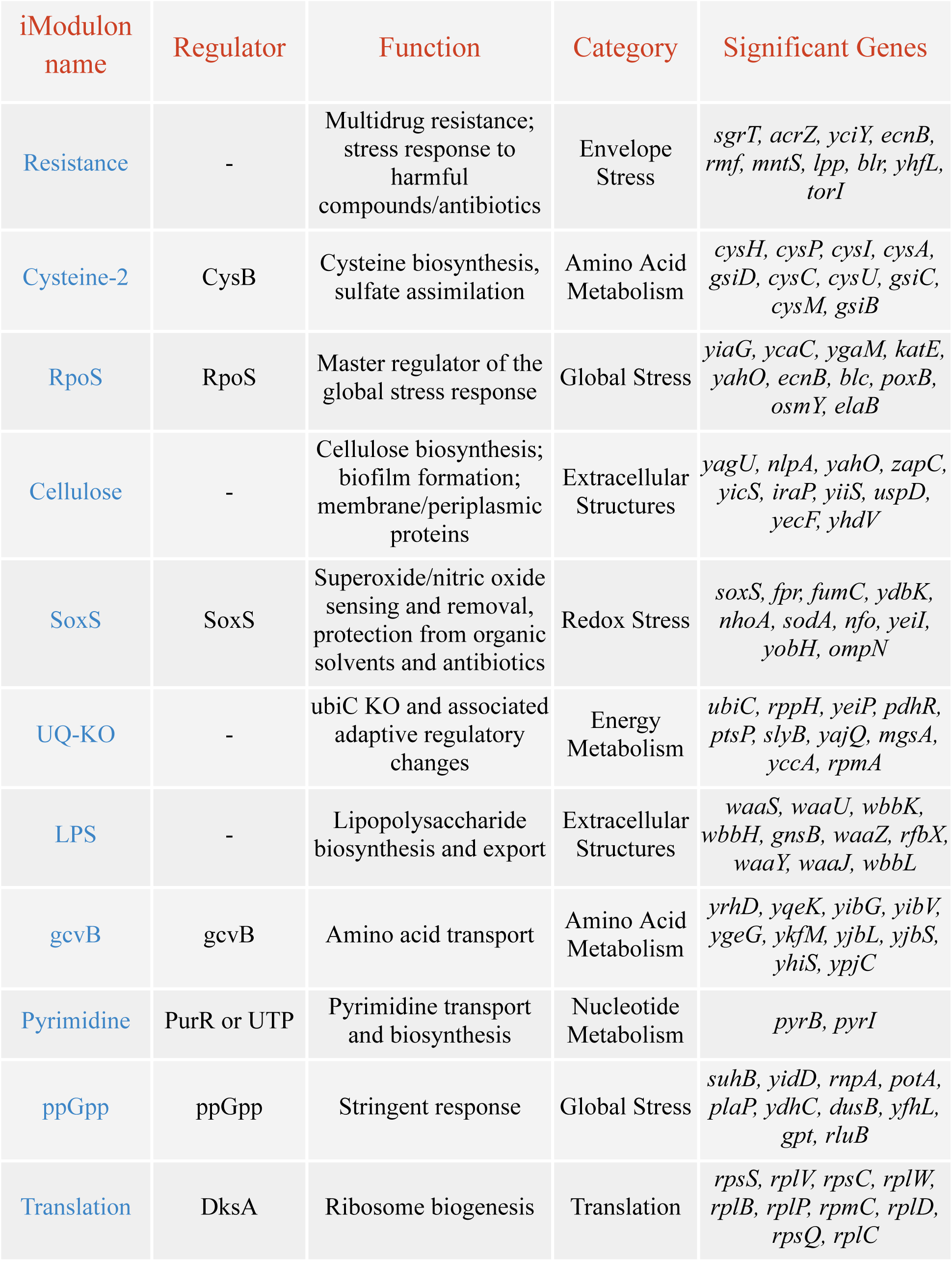
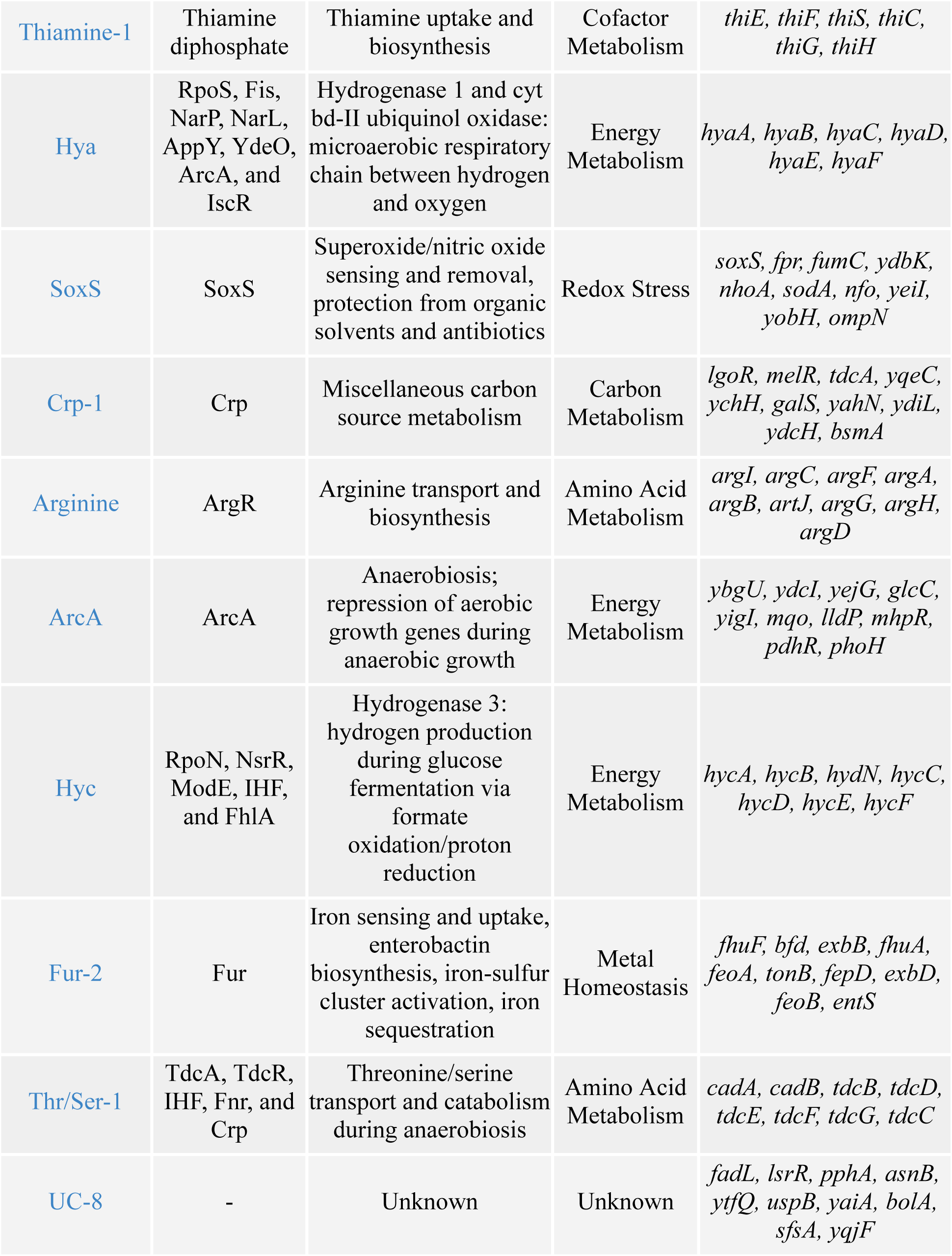
iModulon description.

## Experimental model and subject details

### Escherichia coli BioDesignER

To comprehensively probe the mechanism of the HNQ-mediated EET in *E. coli*, we leveraged the BioDesignER (BioD) strain to make fast-paced genome editing^40^. BioD encodes a refactored λ-Red recombination machinery in its genome, enabling highly efficient recombineering with dsDNA and ssDNA. The λ-Red machinery is encoded under an anhydrotetracycline (aTc) inducible TetR-P_tetO2_ transcriptional control.

### Cell culturing conditions

*E. coli* strains subjected to genome editing were cultured in LB. The *E. coli* strains investigated for EET performances were cultured, washed, and tested in the M9P buffered conditions. The M9P buffer (pH 6.5) comprises 42 mM Na_2_HPO_4_.7H_2_O, 24 mM KH_2_PO_4_, and 9 mM NaCl. A pH of 6.5 was chosen since *E. coli* showed higher HNQ-mediated EET performance in the BES (Figure M1A). The cell cultures were grown with aerobic shaking at 250 rpm at either 37°C or 30°C. All the cell culture media were sterilized using a vacuum filtration system.

### Preparation of cell biomass to test for HNQ-mediated EET

The cell biomass used to investigate EET performance was cultured in two steps. In step one, the pre-cultures inoculated from glycerol stocks were grown for 10 hours at 37°C in M9P-LB media containing 10 g/L tryptone, 5 g/L yeast extract, and 1 mM MgSO_4_. In step two, these pre-cultures were then diluted by a factor of 100 and transferred to M9P-2xYT media containing 16 g/L tryptone and 10 g/L yeast extract, where they were grown for 12 hours at 30°C to generate the cell biomass. These cell biomass-generating cultures were supplemented with 1 mM MgSO_4_, 1x trace metal mix (T1001) purchased from Teknova, 20 mM glycerol, and additional carbon sources catered to the experimental design. The additional carbon source enabled cells to adapt to the carbon source (electron donor) tested under the EET conditions. The cell biomass was prepared by serial washing in a chilled M9P buffer for two rounds (centrifuged at 4000 rpm at 4°C for 15 minutes, and the supernatant was discarded). Each round of washing volume equals twice the cell culture volume. The washed cells were added to a final OD_600_ of either 0.1 or 0.25 to test their EET performances under anaerobic conditions.

### Rationale - Choices of nitrogen sources under HNQ-mediated EET conditions

To simplify elucidating the mechanism of HNQ-mediated EET, the *E. coli* strains were first probed for EET performance under resting conditions. The resting conditions media comprised only the M9P buffer and 2 mM carbon source. This approach was carefully designed to avoid electron flux originating from multiple carbon/nitrogen assimilation pathways and streamline the efforts in probing redox pools elicited by the carbon source. However, once the predominant mechanism of EET under resting conditions was unveiled (Figure 4A-C), a complete-growth medium was used to investigate the additional routes of HNQ-mediated EET. The growth media buffered with M9P (pH 6.5) contained 5 g/L NZ-Amine, 1 mM MgSO_4_, 0.002% v/v Antifoam 204 (Sigma, A8311), and 20 mM glucose.

### Rationale - Choices of carbon sources (electron donors) under HNQ-mediated EET conditions

Glycerol was the preferred choice of electron donor due to its high degree of reduction^126^, which encourages cells to use EET pathways. Our preliminary experiments found that when cells are pre-cultured in M9P-LB with 20 mM glycerol, they used both GlpD and GldA routes of glycerol oxidation under EET testing conditions (Figures 1C and S1D). However, when cells are precultured in M9P-LB without glycerol, our data suggests that the cells use only the GlpD route of glycerol oxidation under EET testing conditions (Figure M1B). Under these conditions, cells can oxidize only a fraction of the provided glycerol, indicating that HNQ-mediated EET does not oxidize the reduced ubiquinone pool (Figure M1C). To benchmark the ideal conditions for investigating the EET mechanism using chronoamperometry (CA) under resting state, we prioritized two conditions: (1) Cells should completely oxidize the limited amount of 2 mM electron donor in about a day, where we expect a bell curve shaped current density vs. time, and (2) The electron flux should originate from the cytoplasm, preferably via NADH pool, since the NADH pool is connected to the rest of the redox biomolecules in the cell. This strategy was expected to reduce the complexity of probing the mechanism of HNQ-mediated EET. While glucose did not fulfill the former condition, pyruvate as the electron donor satisfied both conditions. Thus, unless otherwise mentioned, all the experiments to investigate the mechanism of HNQ-mediated EET under resting conditions were done with pyruvate. However, glucose was used as the electron/carbon source in the experiments with actively growing cells.

### Rationale - Concentration of HNQ used to stimulate EET in *E. coli*

It is important to note that the redox shuttle concentration will influence the mediated EET mechanism and its associated bioenergetics. In this work, we have exogenously supplemented 20 µM HNQ to stimulate EET. Since this study focused on unraveling the mechanism of redox-shuttle-mediated EET, supplementing HNQ at fixed concentrations helps minimize variations across experiments. The concentration of 20 µM was chosen since it gave maximum EET output and did not affect the cell viability under resting/growth conditions (Figure M1D-F). HNQ (Sigma H46805) was dissolved in 100% DMSO. A primary stock of 20 mM HNQ was prepared freshly for every experiment. Thus, with 20 µM HNQ in the EET probing experiments, the unavoidable 14.08 mM of DMSO was introduced in the HNQ-mediated EET experiments. The additional DMSO is critical since it is a potent respiratory electron acceptor under anaerobic conditions and can compete with HNQ-mediated EET. Moreover, DMSO’s presence could alter the cellular carbon and electron fluxes, influencing our experimental approach to understanding the mechanism of HNQ-mediated energy metabolism. However, the most thoroughly investigated strains in this work: AnoxicNull, FermNull, QRedNull, and EsinkNull have all the anaerobic respiratory pathways deleted, including DMSO.

### Rationale - Transcriptomics analysis of cells harvested from amaranth assay over BES

Two aspects can influence the transcriptomics of cells performing EET: (1) The EET pathway and (2) the Redox potential of the electron sink. In the BES system, the extraction of cells performing EET is challenging for the following reasons: (1) In all our experiments, we have injected cells in the BES to a final OD_600_ of 0.1, which is about 5-10 fold lesser than the conventional practice. The small amount of injected cells is initially planktonic, and as the CA experiment proceeds, cells localize in the anode made up of graphite felt (mesh). The time required to isolate an anode from a BES and harvest cells would be sufficient to expose cells to aerobic conditions. Additionally, it could introduce variance amongst replicates. (2) If we were to isolate EET-performing planktonic cells from the BES early, it would not accurately represent the underlying biological question. Since planktonic cells performing mediated EET in a 110 mL BES would exhibit heterogeneity in terms of rate of EET. Thus, to interrogate transcriptomics-associated questions, we designed a proxy EET assay. This assay was conducted in the anaerobic chamber, where instead of an anode, we used a water-soluble and color-changing redox dye called amaranth as the electron sink (Figure M1G). However, the redox potential of amaranth is not the same as the anode poised at 200 mV vs. Ag/AgCl. Thus, amaranth will influence the transcriptome of the investigated strains differently. The amaranth assay is also done in an anaerobic chamber, where cells are exposed to a different gas mixture (96% N_2_, 2% CO_2_, 2% H_2_) compared to BES (100% N_2_). Considering these factors, while the results of the regulon-level transcriptomics analysis from the amaranth assay may not perfectly align with those from the BES, they remain representative of the BES experimental conditions.

## Methods Details

### Gene deletion in BioD

A two-step selection-counterselection approach was utilized to perform clean gene deletions in the BioDesignER (BioD) strain. In the first recombineering step, a GKX selection cassette replaces nearly all of the CDS of the gene to be deleted. In the second step, the GKX cassette is excised by an ssDNA recombination event to achieve the targeted gene deletion (Figure M2A). All the strains generated in this work thrive under aerobic conditions.

All the PCR amplifications of linear dsDNA were done with either the 2x Phanta Max master mix (Dye Plus, Vazyme, P525) or the 2x Phanta Flash Master Mix (Dye Plus, Vazyme, P510). The gel extractions of dsDNA were performed using the FastPure Gel DNA Extraction Kit (Vazyme).

### dsDNA and ssDNA for homologous recombination

The selection cassette (dsDNA) **GKX** cassette comprises a constitutive expression of an sf**G**fp-**K**anR fusion protein alongside a cumate inducible cell lysis gene (phi**X**174). The GKX cassette is integrated into the genome of *E. coli* strain RE880^40^. The GKX selection donors were synthesized by PCR amplifying the GKX cassette from RE880’s genome with primers containing overhangs of 50 bp homology targeting the gene of interest^40^. The suitable PCR amplified selection donors were gel extracted for high purity. On the other hand, the counterselection donor templates of 100 bp ssDNS oligonucleotides containing the tandemly stitched 50 bp homologies with the targeted gene of interest were obtained from Sigma Aldrich. The ssDNA oligonucleotides were designed to have 5’ phosphorothioate base modifications. In the cases of poor ssDNA recombineering efficiency, counterselections were successfully repeated with dsDNA donors, with at least 500 bp homologies obtained from Twist Biosciences.

### Competent cell preparation and recombineering

BioD and its derivative strains were grown in LB Miller broth overnight. The saturated cultures were diluted 100-fold in LB Miller broth supplemented with aTc (100 ng/µL) to induce the λ-Red recombination machinery. Further, the cells were harvested at 0.4-0.6 OD_600_ and prepared for electroporation by washing the cells once with the same culture volume of chilled water and then twice with chilled 10% glycerol. The cells were finally resuspended in 1000^-1^ fold of the initial culture volume with 10% glycerol. The electroporation-guided transformation was done with 50 µL of cells and 400 ng of selection dsDNA donor. The transformations were recovered for three hours and plated on kanamycin LB Miller agar plates. The transformed colonies were confirmed with sfGFP fluorescence and colony PCR. The confirmed colonies were passaged for the next step of clean deletion with kanamycin supplementation in the growth media. The counterselection was performed as described above with either ssDNA or dsDNA, and the transformation recovery was selected on 100 µM cumate. Successful counterselections were confirmed with the loss of sfGFP fluorescence.

### Confirmation of gene deletion

The knockout strains were confirmed with colony PCRs and functional assays. The key strains were genome sequenced and confirmed. The phenotypic confirmation of respiratory and fermentation mutants was performed with anaerobic growth assays. The cell cultures on different anaerobic electron sinks (Nitrate, DMSO, TMAO, and fumarate) were performed in 96-deep well plates in an anaerobic chamber (Whitley A45 anaerobic workstation). The OD_600_ measurements were done using a microplate reader in the anaerobic chamber (Byonoy, Absorbance 96).

### Genome sequencing and mutation analysis pipeline

Genomic DNA was isolated and purified from cell pellets using an Omega Biotek Mag-Bind Bacterial DNA kit (M2350-01). The cell pellets were first resuspended in 800 uL lysis buffer and transferred to a 2 mL tube containing 0.6 ml dry volume mixed 0.5 mm and 0.1 mm ZR BashingBead lysis matrix (Zymo Research S6012-50). The tubes were processed on an Omni International Bead Ruptor 12 Homogenizer instrument (1 cycle, 20 seconds, 6 meters/second) and then centrifuged for 5 minutes at 10,000g. 200 uL of the supernatants were then used for the DNA preparations, following the manufacturer’s protocol.

Whole genome sequencing libraries were prepared from 2 to 40 ng genomic DNA using an NEBNext® Ultra™ II FS DNA Library Prep Kit for Illumina (E7805L), following the manufacturer’s protocol. Library quality was checked on an Agilent TapeStation and quantified using AccuGreen™ High Sensitivity dsDNA Quantitation Solution (Biotium) with an Invitrogen Qubit 2.0 fluorometer. Libraries were then pooled and sequenced on an Illumina Novaseq instrument.

Breseq was run two ways to predict and verify possible genetic mutations in the adapted FermNull strains compared to the FermNull strain^127^. In the first way, both DNAseq reads from FermNull and adapted FermNull strains were compared to the *E. coli* MG1655 reference genome. Mutation predictions specific to adapted FermNull were then considered as genetic adaptations. In the second method a genome was assembled for the FermNull strain based on its DNAseq reads using the spades assembler and prokka to get an annotated genbank file^128,129^. Reads from the adapted FermNull strain were then compared to the assembled FermNull genome. Both methods showed consistent predicted mutations in the adapted FermNull strains.

### Synthetic control on the expression of native genes

In the GldA+ strain, the 5’ region of the genomic *gldA* CDS was replaced with synthetic transcriptional genetic parts for constitutive expression. A strong RBS was designed using the RBS calculator tool on the De Novo platform^130^. A strong and constitutive P_SH039_ promoter was chosen from the non-repetitive synthetic promoter library^131^. A double terminator DT42 was also included upstream of the promoter P_SH039_ (Figure S1E)^132^. The promoter swap was implemented in the BioD derivative strain using the above-described protocol.

The synthetic expression of HNQ-reductase *nfsB* was encoded under an arabinose inducible promoter (P_ara.AM_)^133^ (Figure S4B). The transcriptional activator (AraC) of P_ara.AM_ is constitutively expressed in the stains derived from BioD. The *nfsB* expressing cassette was ordered from Twist Biosciences and integrated into a low-copy part vector using a Golden Gate assembly. The part vector comprises a chloramphenicol-resistant gene and pSC101 origin.

### Testing HNQ-mediated EET using bioelectrochemical systems (BES)

A BES reactor comprises two chambers and a three-electrode configuration (Figure M2B). The two chambers, cathodic and anodic, are filled with 110 mL of M9P buffer/media and separated by a cation exchange membrane (CMI-7000, Membranes International). The cathodic chamber consists of a counter electrode with a 0.5 mm radius of titanium wire (Alfa Aesar). The anodic chamber consists of the working electrode made of titanium wire, which is extended onto a graphite felt with a geometric surface area of 16 cm^2^ (Alfa Aesar), serving as an anode compatible with *E. coli* cells. The anodic chamber is maintained in anaerobic conditions by continuous sparging of N_2_ gas and is also held under constant stirring conditions of 220 rpm by magnetic stir bars (IKA RO10 Magnetic Stirrers). The anodic chamber is also interfaced with an Ag/AgCl reference electrode (CH111, CH Instruments)) saturated with 3 M KCl. Six such water-jacketed BES reactors are connected in series to a temperature-controlled water bath (Adams & Chittenden Scientific Glass; ECO E 4S, Lauda Brinkmann). The temperature of the BES reactors was maintained at 30°C throughout the experiment. Each BES reactor was connected to a potentiostat (Bio-Logic Science Instruments, TN, USA, model - VSP300) to perform electrochemical measurements.

All the electrochemical measurements were done with the anode poised at 200 mV vs. Ag/AgCl at the potential resolution of 100 µV. In the cyclic chronoamperometry (CA) measurements, the current levels were recorded every 36 seconds. In the cyclic voltammetry (CV) measurements, the scan rate of 1 mV/s was used over the potential window of -500 mv to 0 mV. The *E. coli* strains under investigation were added roughly 12-14 hours of nitrogen gas spurging in the BESs. This timeframe ensures the complete elimination of oxygen from the BES setup and stabilizes current baseline levels. The EET enabling components, consisting of cells, HNQ, and the electron donor, were added in this order. This sequence was chosen to facilitate the oxidation of reducing equivalents accumulated from the prior aerobic growth. This approach allowed us to evaluate the EET output in response to the electron donor oxidized in the anaerobic BES conditions.

The EET output from CA experiments is interpreted in three ways: (1) Peak current density (mA/m^2^; Ampere, meter) - The highest current value corresponding to a state where the highest fraction of the provided HNQ is reduced (2) Stead state current density (mA/m^2^) - The current value at the end of the CA run (3) Charge (Q; Coulombs) - The total area under CA plot (current vs. time) corresponding to the total charge deposited on the anode by the end of the CA run. All these values are generated in the software (Bio-Logic, EC-lab) that operates the potentiostat.

As the CA experiments progress, the cells localize in the graphite felt. Thus, to measure the final OD_600_, the BES is disconnected from the potentiostat and the temperature-controlled water bath system, followed by vigorous shaking to dislodge the cells. The cell OD_600_ was then measured on a spectrophotometer (Agilent, Cary Series UV-Vis).

### Testing HNQ-mediated EET using a colorimetric assay

The colorimetric assay utilized amaranth, an azo dye (Sigma A1016), as the electron sink. This colorimetric assay was initially validated with an assay media containing 125 µM amaranth in the M9P complete-growth medium with 20 mM pyruvate. In this assay, cells were inoculated at OD_600_ of 0.25 and incubated in a shaker (IKA, VXR basic Vibrax) in the anaerobic chamber for 12 hours. However, the cells exposed to amaranth at 125 µM for a long time were found to have RNA degradation, especially in the *E. coli* strains (FermNull and EsinkNull) with impaired anaerobic growth. Thus, the assay was redesigned to use 12.5 µM amaranth dye with shaking under anaerobic conditions for 2 hours. This redesigned assay was used to collect cells for transcriptomics analysis. The degree of amaranth reduction was measured by probing absorbance at 518 nm (Abs_518_) of the spent assay media using a spectrophotometer (Agilent, Cary Series UV-Vis). The reported percentage amaranth reduction was calculated as follows:

Amaranth reduction (%) = [(No cell control)_Abs518_ - (*E. coli* strain)_Abs518_]/(No cell control)_Abs518_

Note: To validate if the amaranth reduction assay^88,90^ mimics the EET phenotype shown by the strains in BES experiments, we probed the AnoxicNull, QRedNull, and EsinkNull strains (Figure 5A). The amaranth reduction was measured by quantifying the decreased absorbance of the spent media at 518 nm. The AnoxicNull, QRedNull, and EsinkNull show amaranth reduction of 59.7%, 30.7%, and 15.9%, respectively (Figure M1H). This data indicates EsinkNull has the lowest HNQ reduction ability and is consistent with the EET assessments in BES. In this assay, all the tested strains consumed similar pyruvate levels (Figure M1I); however, EsinkNull retained ∼30-fold higher formate concentrations (Figure M1J). The retention of formate is attributed to the formate dehydrogenase deletions in EsinkNull. EsinkNull also generated ∼1.7-fold higher acetate levels (Figure M1K), suggesting that AnoxicNull and QRedNull have relatively lower carbon flux through ATP-generating acetyl-CoA/acetate node. These data indicate that the amaranth can serve as a proxy for the EET tests in BES, and the cells could be effortlessly harvested for transcriptomics analysis.

### Measuring metabolite concentrations using HPLC

Quantifications of glycerol, glucose, and other organic acids were done using HPLC (Shimadzu, Kyoto, Japan). An isocratic mobile phase of 30 mM H_2_SO_4_ was operated on the Aminex Organic Acid Analysis column (Bio-Rad, HPX-87H 300 × 7.8 mm) with a flow rate of 0.3 L min^−1^. The Aminex column was held at 60°C, and the compounds of interest were separated with discernible peaks. Glycerol and glucose were measured using a refractive index detector (Shimadzu, RID-20A, 120V), whereas the organic acids were measured using a UV detector (Shimadzu, SPD-M20A). No gaseous products were measured in this study.

### Modeling of HNQ-mediated EET

We performed Flux Balance Analysis using the latest genome-scale metabolic model of *E. coli,* iML1515^134^. We generated three sets of strains for analysis: 1) wild type, which is identical to the base iML1515 model; 2) EsinkNull and FermNull, which have the sets of redox reaction knockouts described in this manuscript (see Strain list); and 3) EsinkNull (and FermNull) + nfsB, which has the EsinkNull (and FermNull) reaction sets knocked out but the addition of the reactions for naphthoquinone reduction by NADH (NADH + Oxidized HNQ ↔ NAD + Reduced HNQ), and regeneration by the anode (Reduced HNQ ↔ Oxidized HNQ). FBA was performed in a standard manner^95^ using the COBRA Toolbox^135^ in Matlab with the Gurobi solver version 9.1.1. We utilized growth as the primary objective for simulations, with glucose uptake constrained to 10 mmol/gDW/hr, oxygen constrained to 0 mmol/gDW/hr (anaerobic), with a secondary quadratic objective to minimize the Euclidean norm of the flux state while constrained to the maximal growth rate. Additionally, the NAD(P) transhydrogenase flux was capped to predicted wild-type levels of 2 mmol/gDW/hr in all simulations to prevent the use of futile redox cycles to oxidize NADH freely. Genes were knocked out respective of logical relationships in reaction gene-to-protein relationships (GPRs), such that a reaction flux was constrained to 0 mmol/gDW/hr if and only if all of its possible catalyzing protein complexes have at least one essential gene knocked out. Simulations were performed in Matlab using the COBRA toolbox^135^.

We then analyzed the proteome allocation using the iJL1678b Metabolism and Macromolecular Expression (ME) model^98^. We generated three strains for analysis: 1) wild type, identical to the base iJL1678b model; 2) FermNull, as described above; and 3) FermNull + nfsB, as described above. We estimated the NfsB k_eff_ value to be 57.4 in the model using the transcriptome-constrained wild-type flux state and experimental absolute proteomics from this study^136^. All strains were grown anaerobically, with growth being the primary objective for simulations. The Wild Type strain was simulated under two conditions: 1) Fermentation, which was grown anaerobically with glucose uptake constrained to 10 mmol/gDW/hr, and 2) Nitrate Respiration, which was grown similarly but additionally with nitrate uptake constrained to 30 mmol/gDW/hr. The FermNull and FermNull + nfsB strains were constrained to the same growth rates calculated from the iML1515 simulations. Proteome allocation was analyzed using the simulation results for ETC, Glycolysis, and Fermentation proteins. Proteome allocation related to stress iModulons (SoxS, OxyR, ppGpp, CpxR, and RpoS) were analyzed for the FermNull strains by calculating the proteome allocated to the genes enriched in each iModulons. Simulations were performed using the COBRAme Python software package^98^.

### RNA-seq and transcriptomics analysis pipeline

6 ml of cell culture (OD_600_ ∼ 0.25) was immediately added to two volumes of RNA protection solution (RNAprotect Bacteria Reagent, Qiagen 76506), vortexed for 5 s, incubated at room temperature for 5 min, and immediately centrifuged for 15 min at 4000 rpm. The supernatant was decanted, and the cell pellet was stored at the −80 °C. Cell pellets were thawed, and total RNA was prepared using a PuroMAG™ Total RNA Purification Kit (Luna Nanotech Inc., catalog number NKM051-96), following the vendor’s manual, including a 15-minute DNase treatment at room temperature. RNA was quantified using a Nanodrop, and quality was assessed by running on an Agilent TapeStation. The rRNA was removed using the RiboRid procedure with oligonucleotide probes specific to *E. coli* ribosomal RNA. Following the manufacturer’s protocol, a KAPA RNA HyperPrep kit (catalog number 08098107702) was used to create sequencing libraries with an average insert length of around ∼300 bp. Libraries were run on an AVITI instrument (Element Biosciences).

RNAseq abundance computation, quality checks, processing and Independent component analysis (ICA) for calculating iModulons were done using the modulome workflow^137^. Differential gene expression was done using Deseq enabled with the pydeseq library in python. An adjusted P_value_ of 0.05 and log_2_(FoldChange) of 1.5 was taken as the threshold for determining significant genes. For figure 5, due to a new ICA capturing strain level variance as signals, we inferred the iModulon activities using the existing *E. coli* structure by deleting all the genetic knock outs across the strains. This resulted in capturing within strain level differences across HNQ and DMSO better while preserving variance across strains. For figure 6, we ran ICA on all data up to figure 5. We then used this newly obtained iModulon structure to infer activities for data in figure 6. This was done to see what the newly obtained iModulons with figure 5 data could tell us about unseen/figure 6 data and how iModulon activities changed across the datasets.

Note: In some experiments, we compared the transcriptome of the strains under no-EET conditions. Here, we aimed to investigate the transcriptome makeup of the cells as they were introduced into the BES under EET conditions. Chronoamperometry experiments indicate that the EET machinery reducing HNQ in the respective strains (BioD, FermNull, and adapted FermNulls) was already present when these strains were introduced into the BES.

---

**Supplementary Figure 1.**
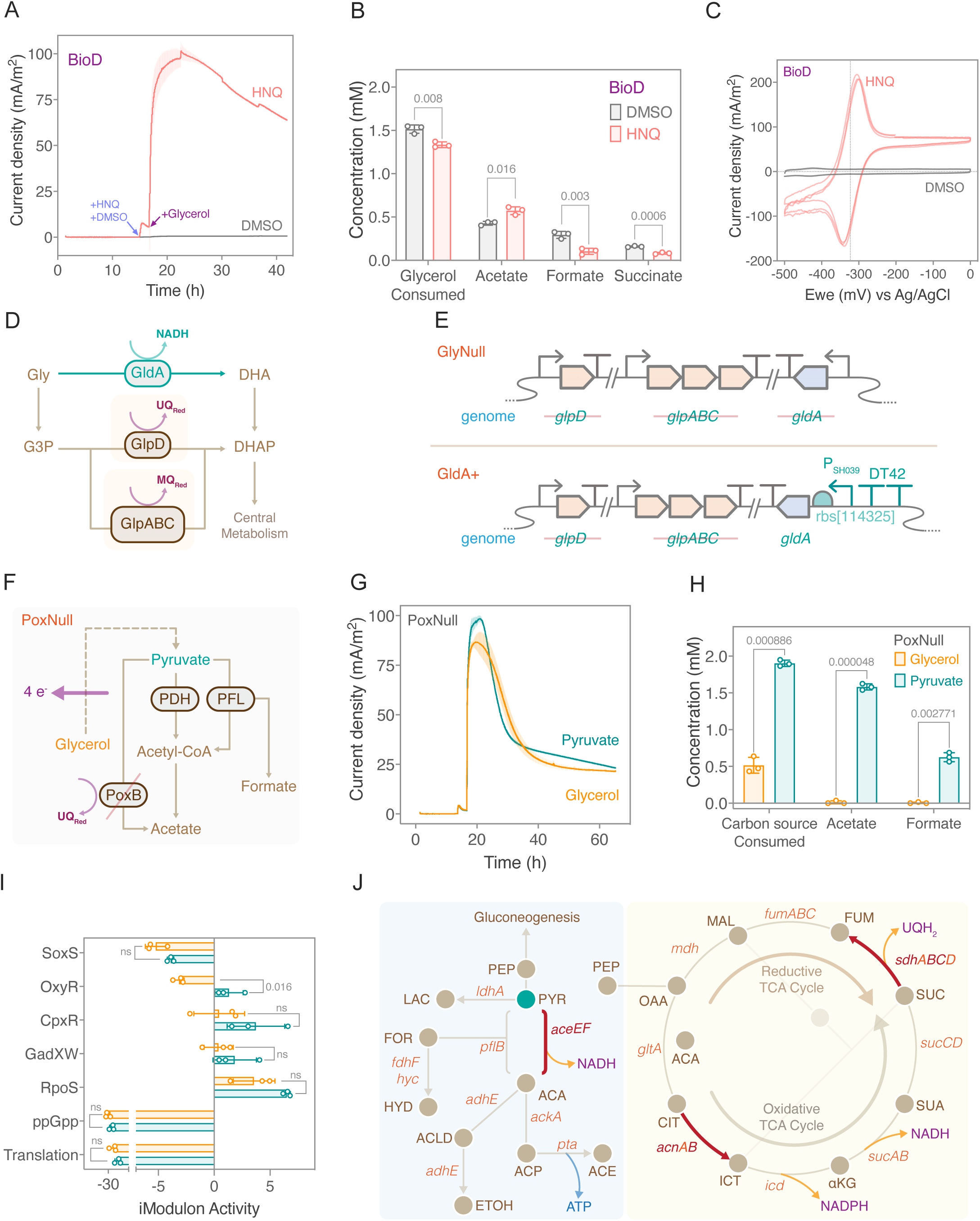
related to Figure 1. (A) Chronoamperometry of BioD with and without HNQ under resting conditions, showing that HNQ stimulates EET upon the addition of glycerol as the electron donor. (B) Glycerol consumption and the metabolite profile of the spent media, showing that cells under both conditions are metabolically active. The data in (A) and (B) represent mean ± standard deviation obtained in triplicate BES experiments. P-values are calculated using the Welch’s t-test with a significance threshold of 0.05. (C) Cyclic voltammogram (CV) of BioD with HNQ and DMSO, showing that HNQ serves as a redox shuttle between metabolically active cells and the anode. The CV waveforms are plotted from three independent BES experiments. (D) *E. coli* consumes glycerol via three routes: GlpD and GlpABC are glycerol 3-phosphate dehydrogenases that relay electrons to the quinone pool, and GldA is a glycerol dehydrogenase that relays electrons to the NADH pool. GlpD is the preferred route under aerobic conditions, whereas GlpABC is the preferred route of anaerobic glycerol oxidation. GldA expression, on the other hand, is independent of aerobicity. (E) Genotypic representation of GlyNull and GldA+ strains, showing that GlyNull is devoid of all the three routes of glycerol consumption whereas GldA+ has intact *gldA*, under a synthetic constitutive promoter, as the sole route of glycerol consumption. (F) A schematic showing the catabolism of glycerol and pyruvate as the electron donors in the PoxNull strain. Since glycerol is more reduced than pyruvate, it can release 4 extra electrons. So, a higher EET output is expected on the complete oxidation of glycerol compared to pyruvate. (G) PoxNull generates similar EET outputs with either glycerol or pyruvate as electron donors under resting conditions. The strains in this experiment were supplemented with 20 mM glycerol and 20 mM pyruvate in both pre-culture and biomass-generating cultures. (H) Carbon source consumption and the corresponding metabolite profile of the spent media, showing that PoxNull consumed all of the pyruvate (∼2mM), whereas it consumed only about 25% of the total glycerol provided. The data in (G) and (H) represent mean ± standard deviation obtained in triplicate BES experiments. P-values are calculated using the Welch’s t-test with a significance threshold of 0.05. (I) Activity levels of stress-related imodulons, showing that only OxyR activity difference is statistically significant between the samples, and no other stresses are reflected in the transcriptome. P-values are calculated using the Welch’s t-test with a significance threshold of 0.05. (J) Higher expression levels of NADH-generating pathways are observed with HNQ. Highly expressed genes include *aceE* and *aceF*, subunits of pyruvate dehydrogenase, which release NADH. Additionally, the *acnB* subunit of aconitate hydratase and the *sdhB* and *sdhC* subunits of succinate dehydrogenase, which are part of the oxidative TCA cycle, are highly expressed.

**Supplementary Figure 2.**
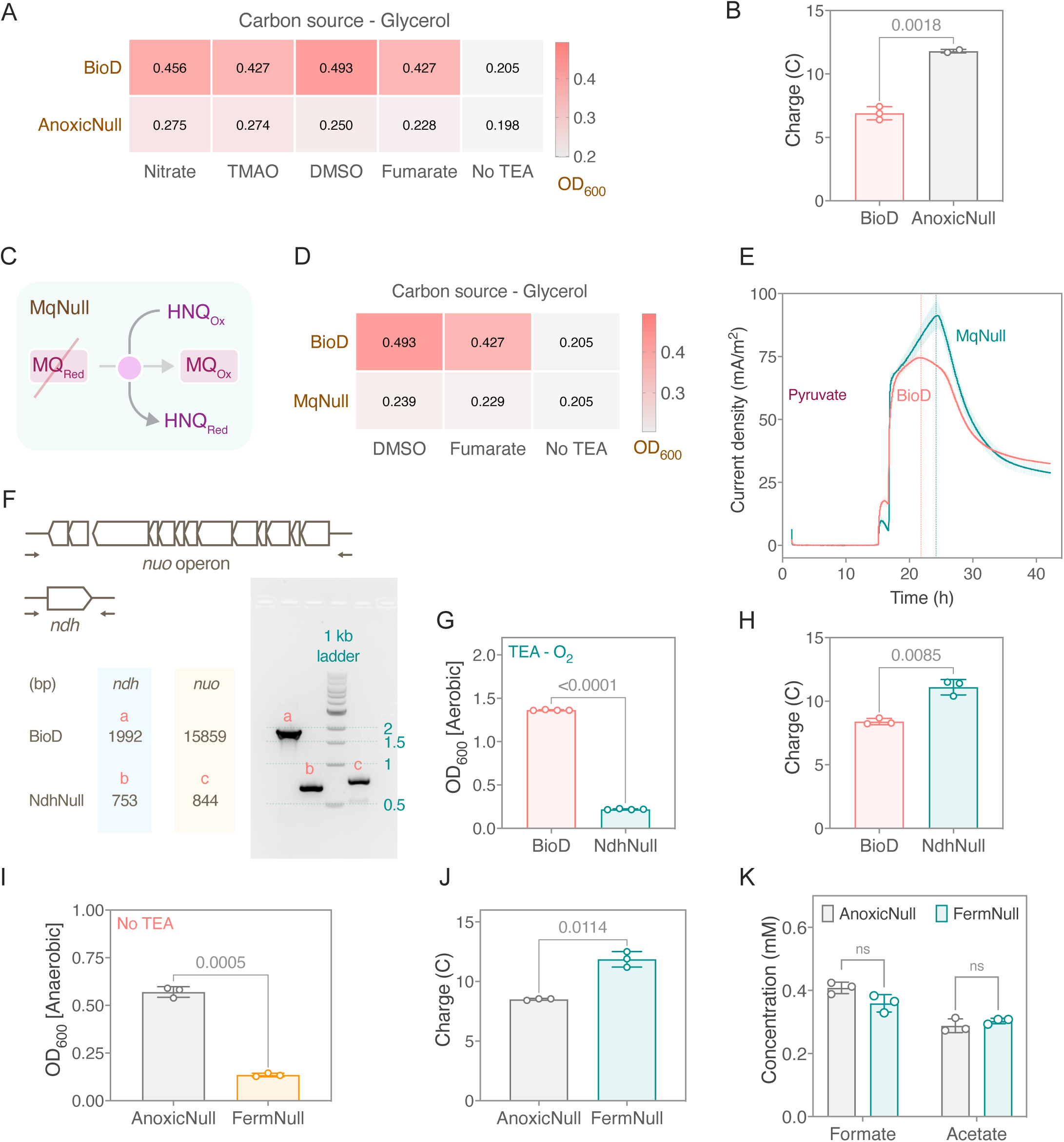
related to Figure 2 (A) OD_600_ of BioD and AnoxicNull strains after 30 hours of anaerobic growth on various respiratory electron acceptors in M9P-2xYT media, showing that AnoxicNull consistently achieves lower growth levels than BioD in the presence of TEAs but has similar growth levels as BioD with no electron acceptor conditions. This data confirms that nitrate, TMAO, DMSO, and fumarate respiratory pathways are successfully eliminated in the AnoxicNull strain. (B) The total charge deposited by BioD and AnoxicNull on the anode with 2 mM pyruvate as the electron donor, showing that AnoxicNull yields higher electron transfer onto the anode than BioD. (C) A schematic showing the menaquinone pool that can reduce HNQ is deleted in the MqNull strain (D) OD_600_ of BioD and MqNull strains after 30 hours of anaerobic growth on DMSO, fumarate, and no TEA conditions in M9P-2xYT media. The lack of a menaquinone pool prevents DMSO and fumarate from being used in anaerobic respiration as TEA, confirming that menaquinone biosynthesis is successfully knocked out in the MqNull strain. (E) Chronoamperometry of the BioD and MqNull strains under resting conditions, showing that the peak current density of MqNull is higher than that in BioD. This data suggests that the menaquinone pool is not involved in reducing HNQ under EET conditions. The data represent mean ± standard deviation obtained in duplicate BES experiments. (F) Colony PCR confirmation of the *ndh* gene and the *nuo* operon knockout in NdhNull. The *nuo* operon size in BioD is very large to amplify and distinctly show on the agarose gel; thus, the *nuo* locus amplification is shown only in the strain with *nuo* operon knockout. (G) OD_600_ of BioD and NdhNull after 10 hours of aerobic growth in M9P-LB media, showing that lack of NADH dehydrogenases abates biomass growth. This growth assay phenotypically validates the genotype of NdhNull. (H) The total charge deposited by BioD and NdhNull on the anode with 2 mM pyruvate as the electron donor, showing that NdhNull facilitates higher electron transfer onto the anode than BioD. (I) Anaerobic growth of AnoxicNull and FermNull in M9P-LB media, showing that the lack of fermentation pathways reduces biomass growth of the FermNull strain. This anaerobic growth assay phenotypically confirms the deletion of fermentative pathways in the FermNull strain. The data represent mean ± standard deviation obtained in three biological replicates. (J) The total charge deposited by AnoxicNull and FermNull on the anode with 2 mM glucose as the electron donor, showing that FermNull yields higher electron transfer onto the anode than AnoxicNull. (K) The metabolite profile of the spent BES media, showing that there is no difference between the amount of formate and acetate generated by the AnoxicNull and FermNull strains. All the P-values are calculated using the Welch’s t-test with a significance threshold of 0.05.

**Supplementary Figure 3.**
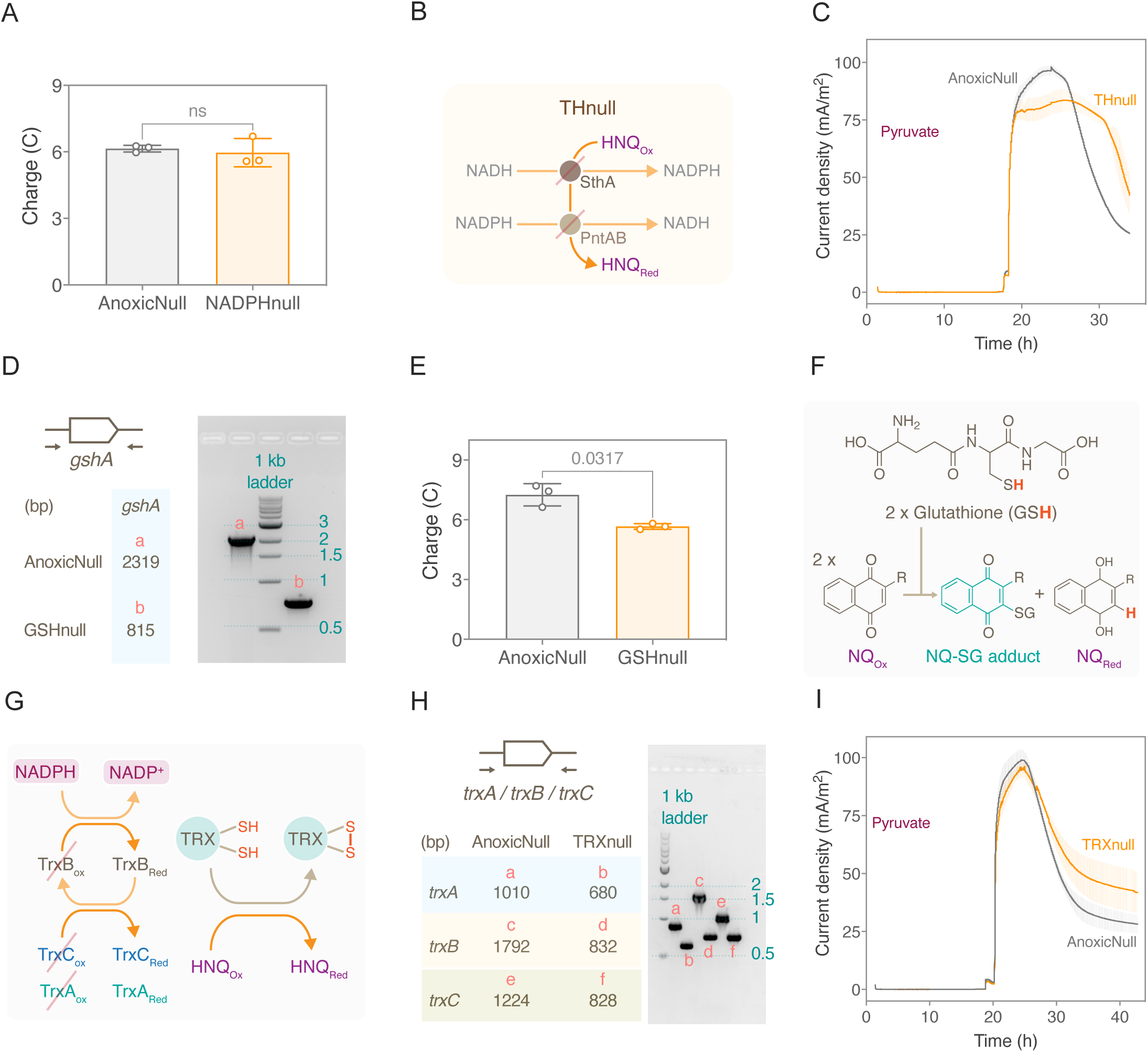
related to Figure 3 (A) The total charge deposited by AnoxicNull and NADPHnull on the anode with 2 mM pyruvate as the electron donor, showing that both strains deposit similar amounts of electrons on the anode. The P-value is calculated using Welch’s t-test. (B) Transhydrogenases that can reduce HNQ are deleted in the Thnull strain (C) Chronoamperometry of AnoxicNull and THnull strains under resting conditions, showing no visible differences in the EET performance. This result indicates that the transhydrogenases do not reduce HNQ. (D) Colony PCR confirmation of the *gshA* gene knockout in GSHnull (E) The total charge deposited by AnoxicNull and GSHnull on the anode with 2 mM pyruvate as the electron donor, showing that GSHnull yields lower electron transfer onto the anode compared to AnoxicNull. The P-value is calculated using the Welch’s t-test with a significance threshold of 0.05.(F) Reduced glutathione can abiotically react with oxidized naphthoquinones (NQ) to form an NQ-SG adduct and a reduced version of the NQ. (G) The NADPH pool maintains the reduced form of the thioredoxin (TRX), which can transfer electrons to HNQ. (H) Colony PCR confirmation of the *trxA, trxB, and trxC* gene knockouts in TRXnull (I) Chronoamperometry of AnoxicNull and TRXnull strains under resting conditions, showing no visible differences in the EET performance. This result indicates that the thioredoxin pool does not reduce HNQ in the conditions tested. The data represent mean ± standard deviation obtained in triplicate BES experiments.

**Supplementary Figure 4.**
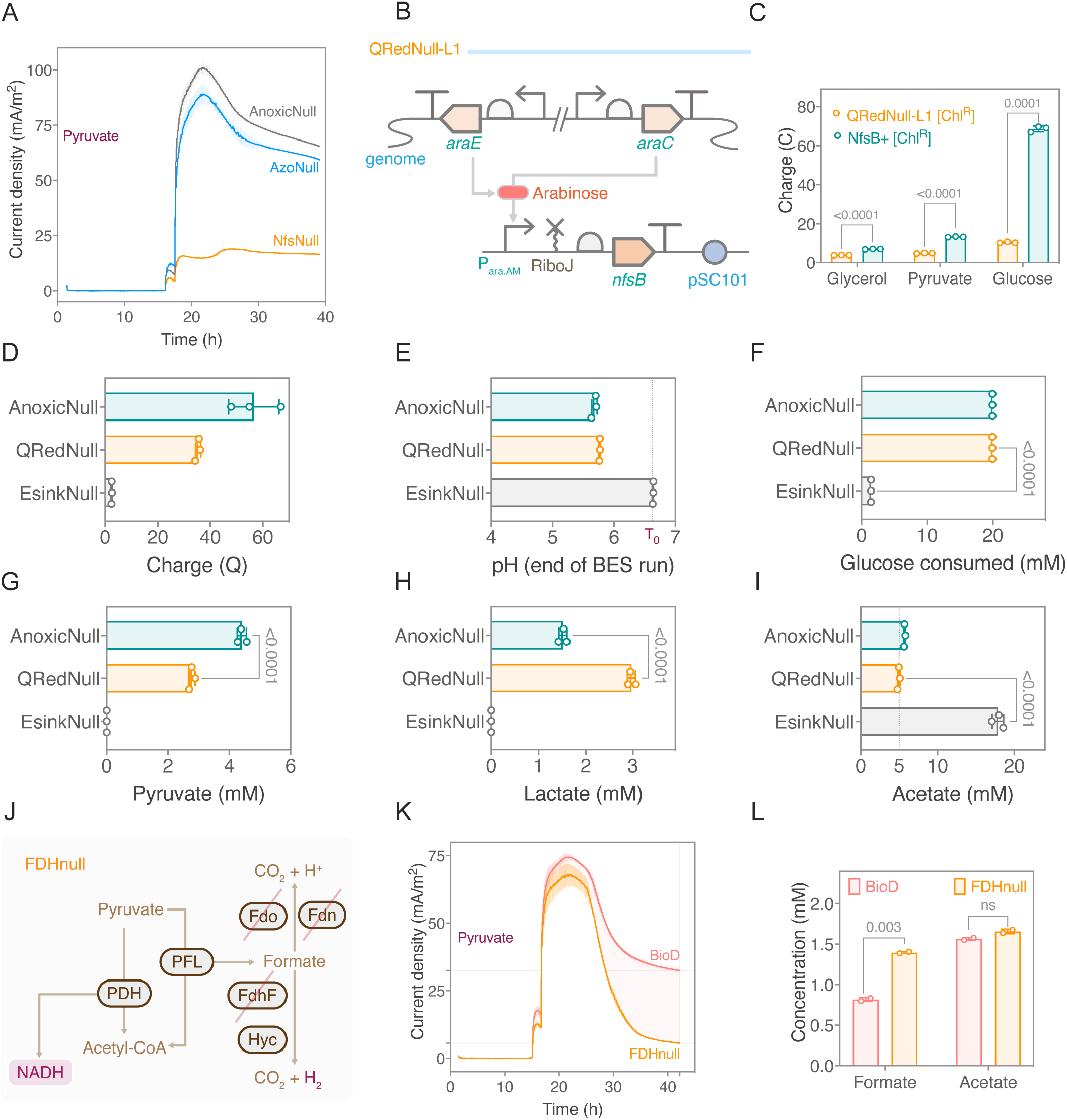
related to Figure 4 (A) Chronoamperometry of the AnoxicNull, the AzoNull, and the NfsNull strains under resting conditions. The EET levels in NfsNull are substantially diminished compared to the AnoxicNull and AzoNull strains. This data indicates that nitroreductase NfsA and NfsB can reduce HNQ. The data represent mean ± standard deviation obtained in duplicate BES experiments. (B) A schematic showing the plasmid expression design of NfsB in the QRedNull-L1 strain. (C) The total charge deposited by QRedNull-L1 and NfsB+ on the anode with different electron donors, showing that complementing NfsB on a plasmid in the QRedNull-L1 strain enables higher charge deposition on the anode. (D) The total charge deposited by BioD, AnoxicNull, QRedNull, and EsinkNull on the anode under growing conditions, demonstrating that the amount of charge deposited on the anode is directly proportional to the strains’ ability to grow using fermentative pathways. (E) pH measured at the end of the BES run, suggesting that EsinkNull did not significantly oxidize glucose. (F) Glucose consumed by the end of the BES run suggesting that EsinkNull is limited in oxidizing glucose due to a lack of available electron sink. The (G) pyruvate, (H) lactate, and (I) acetate profiles of the spent BES media of BioD, AnoxicNull, QRedNull, and EsinkNull, suggesting that all strains are metabolically active. (J) A schematic depicting that all three formate dehydrogenases are deleted in the FDHnull strain. (K) Chronoamperometry of the BioD and FDHnull strains under resting conditions, showing that BioD maintains a higher steady current density level than the FDHnull strain. This data indicates the flow of electrons in the formate catabolism contributes to HNQ-mediated EET. (L) The metabolite profile of the spent BES media, showing that there are different levels of formate retention in the BioD and the FDHnull strains. This data indicates that BioD performs additional EET at the expense of formate. The data in (K) and (L) represent mean ± standard deviation obtained in duplicate BES experiments. All the P-values are calculated using Welch’s t-test, with a significance threshold of 0.05.

**Supplementary Figure 5.**
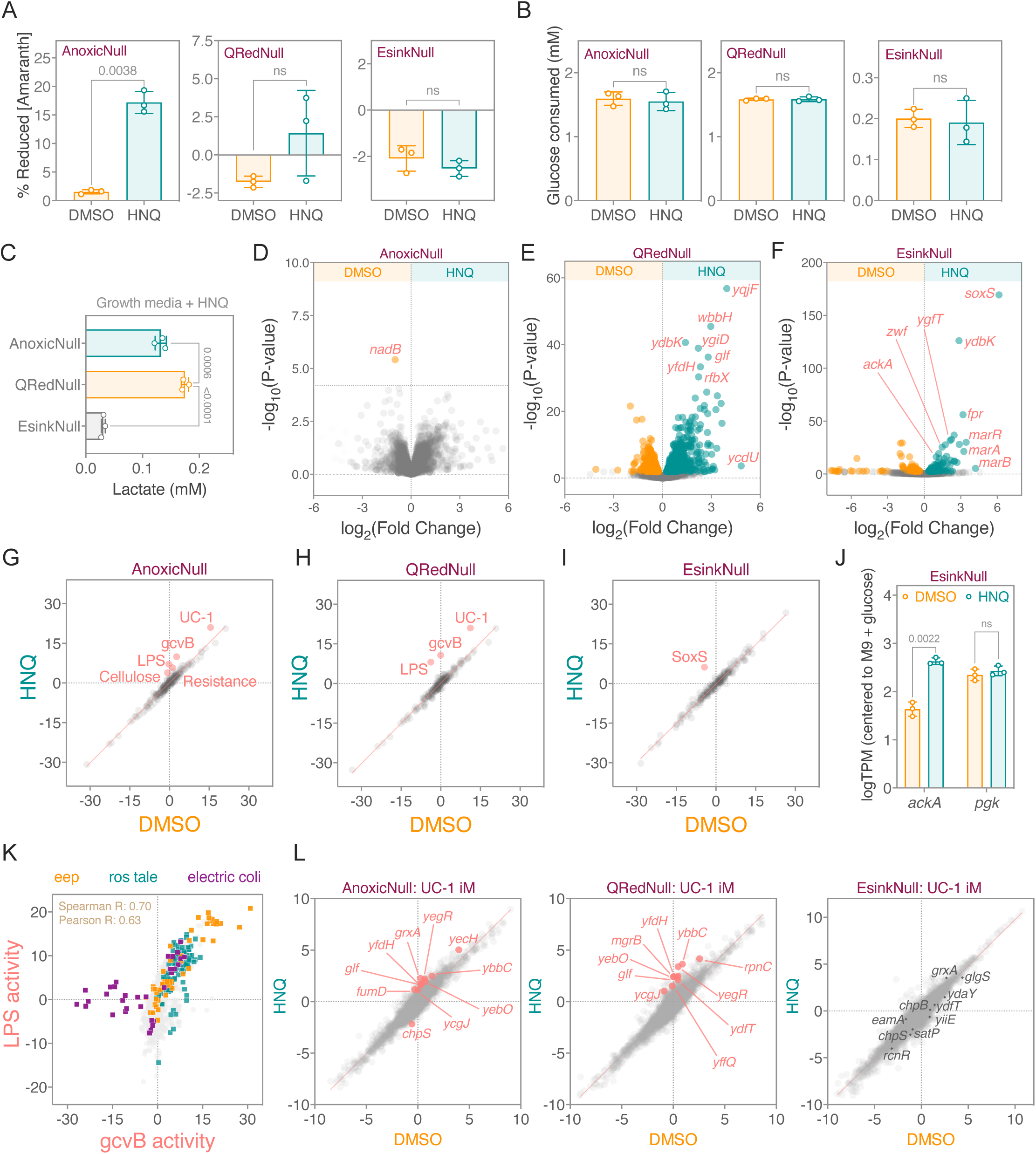
related to Figure 5 (A) The percentage reduction of amaranth by AnoxicNull, QRedNull, and EsinkNull in the presence of DMSO and HNQ, showing that, using HNQ, the AnoxicNull strain significantly reduces amaranth, whereas QRedNull and EsinkNull barely perform measurable EET in 1.5 hours. (B) Glucose consumed in the amaranth assay of AnoxicNull, QRedNull, and EsinkNull at the end of 1.5 hours, showing that both AnoxicNull and QRedNull consume glucose, whereas EsinkNull does not. This data indicates the presence of either fermentative pathways or HNQ-mediated EET in AnoxicNull and QRedNull allows oxidation of glucose. The P-values in (A) and (B) are calculated using Welch’s t-test, with a significance threshold of 0.05. (C) Lactate measured in the amaranth assay of AnoxicNull, QRedNull, and EsinkNull at the end of 1.5 hours, showing that QRedNull produces a higher lactate than AnoxicNull and EsinkNull. This result validates the genetic background of the strains and matches with the lactate production trend previously seen in the BES condition. The P-values are calculated using one-way ANOVA, with a significance threshold of 0.05. The data in (A) to (C) represents mean ± standard deviation obtained in triplicate experiments. (D) Differential gene expression analysis (DEG) of AnoxicNull in DMSO alone and with HNQ in amaranth assay, showing down-regulation of the NAD biosynthesis gene *nadB* when HNQ is supplemented. (E) Differential gene expression analysis (DEG) of QRedNull in DMSO alone and with HNQ in amaranth assay, showing an HNQ-induced upregulation of several LPS-related genes and other genes known to be activated by quinones including *ygiD* and *yqiF*. (F) Differential gene expression analysis (DEG) of EsinkNull in DMSO alone and with HNQ in amaranth assay, showing that HNQ activates various redox genes including *soxS*, *zwf*, *ydbK*, and *fpr*, as well as antibiotic resistance-associated *mar* genes. (G) Differential iModulon activity (DiMA) for AnoxicNull with and without HNQ supplementation, showing simultaneous HNQ-induced activation of the LPS, gcvB, and UC-1 iModulons as well as cellular and two complex iModulons called cellulose (containing diverse functions) and resistance (associated with multi-drug resistance). (H) Differential iModulon activity (DiMA) for QRedNull with and without HNQ supplementation, showing HNQ-induced activation of the LPS, gcvB, and UC-1 iModulons. (I) Differential iModulon activity (DiMA) for EsinkNull with and without HNQ supplementation, showing that HNQ only differentially induces the SoxS iModulon in this strain. (J) Transcript levels of the ATP-generating genes *ackA* and *pgk* in EsinkNull. P-values are calculated using the Welch’s t-test, with a significance threshold of 0.05. (K) The activity of the LPS and gcvB iModulons across the PRECISE1k database of *E. coli* gene expression. Samples from this project and other projects with high activity of these iModulons are highlighted. (L) Different gene expression plots for AnoxicNull, QRedNull, and ESinkNull, with highly perturbed genes within the Uncharacterized-1 (UC-1) iModulon highlighted. Diverse redox-associated genes (e.g. glf, fumD) and y-genes appear in this component. Gene expression was centered on an aerobic glucose reference condition.

**Supplementary Figure 6.**
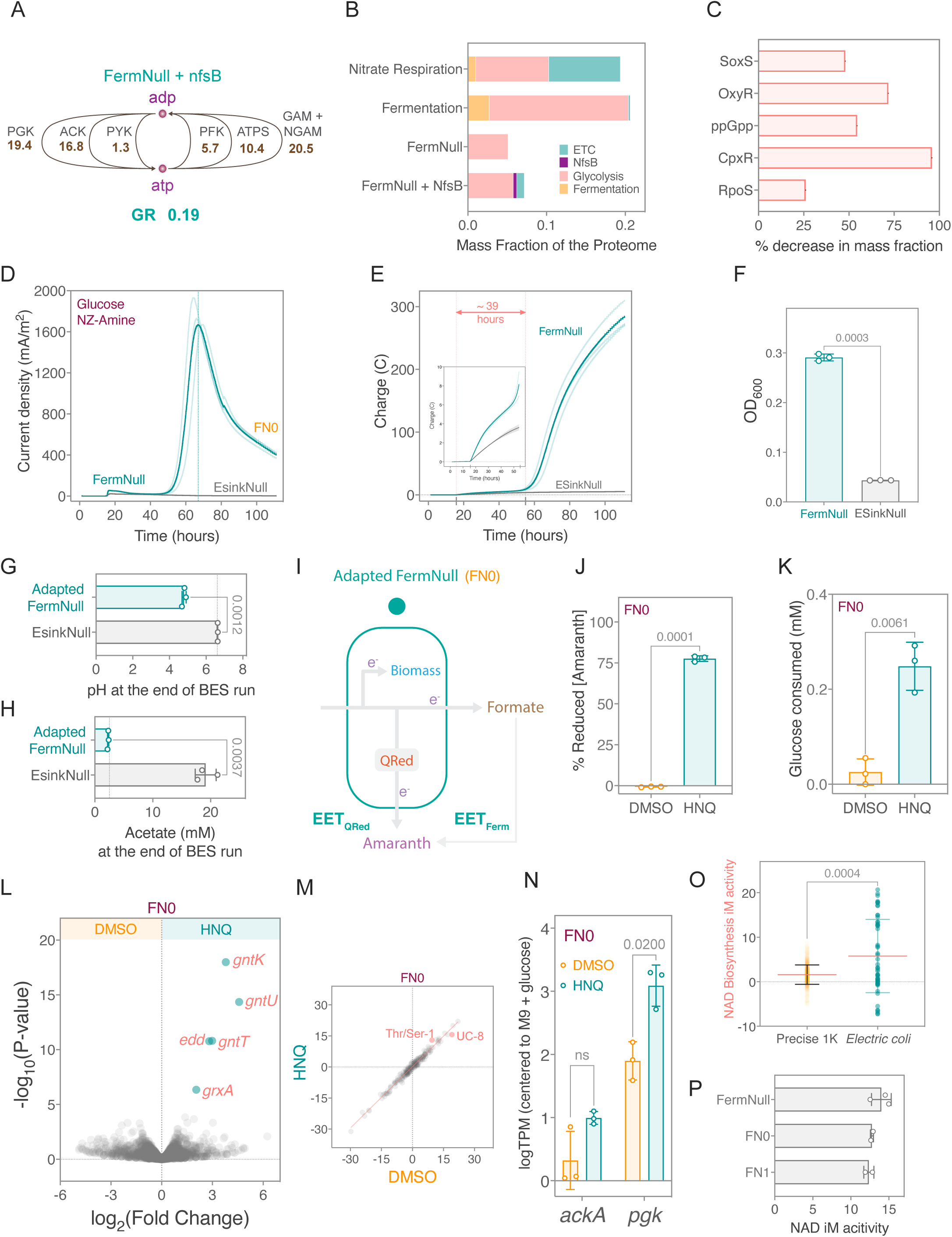
related to Figure 6 (A) Flux balance analysis of the FermNull+nfsB strain under anaerobic growth on glucose shows the ATP production pathways supporting elevated growth. Fluxes in mmol/gDW/hr units, growth rate (GR) in 1/hr units. The listed fluxes represent >95% of the turnover of ATP/ADP. Reaction abbreviations - PGK: Phosphoglycerate Kinase. ACK: Acetate Kinase. PYK: Pyruvate Kinase. PFK: Phosphofructokinase. ATPS: ATP Synthase. GAM: Growth-associated Maintenance. NGAM: Non-growth-associated Maintenance (B) A stacked bar chart that displays the mass fraction of the proteome allocated to ETC, Glycolysis, Fermentation sectors, as well as NfsB. This is done for a wild-type strain undergoing nitrate respiration, fermentation only, and the FermNull strain with and without the addition of NfsB. (C) A bar chart that shows the percent decrease in the mass fraction of the proteome allocated to five stress-related iModulons upon the addition of a QRed (NfsB) to FermNull. (D) Chronoamperometry of the FermNull and the EsinkNull strains under growth conditions, showing that FermNull transforms into generating a high EET output after several hours in the BES. This indicates that the FermNull strain is undergoing changes that increase the flux through the HNQ-mediated EET pathway. (E) The total charge deposited on the anode by FermNull and EsinkNull over time, showing that FermNull consistently deposits a higher charge value over time than EsinkNull. (F) OD_600_ of the FermNull and the EsinkNull strains post BES run under growth conditions, showing FermNull grew to higher biomass than EsinkNull. This indicates that HNQ-mediated EET can enable growth under anaerobic conditions. The data in (D), (E), and (F) represent mean ± standard deviation obtained in triplicate BES experiments. (G) pH measured at the end of the BES run, suggesting that as compared to EsinkNull, adapted-FermNull has conducted higher catabolism and EET. (H) Acetate produced by the end of the BES run, showing that EsinkNull has a higher carbon-flux going into acetate biosynthesis than the adapted-FermNul strain. The data represented in (G) and (H) are the means ± standard deviations obtained in triplicate BES experiments. (I) The FN0 strain derived from FermNull cannot grow on fermentative pathways but performs quinone reductase facilitated HNQ-mediated EET. (J) The percentage reduction of amaranth by FN0, showing that in the presence of HNQ, FN0 can significantly reduce amaranth in 1.5 hours. This result, along with the Sup fig 5A, corroborates with the EET phenotypes seen in the BES set with FN0 and EsinkNull. (K) Glucose consumed in the amaranth assay of FN0 at the end of 1.5 hours, showing that FN0 significantly oxidizes glucose only in the presence of HNQ. (L) Differential gene expression (DEG) analysis of FN0 with and without HNQ supplementation, showing an activation of gluconate degradation genes *gntKUT* and *edd* and the glutaredoxin gene *grxA*. (M) Differential iModulon activity (DiMA) for FN0 with and without HNQ supplementation, showing an up-regulation of an amino acid iModulon (Thr/Ser-1) and a down-regulation of an iModulon containing poorly characterized genes (UC-8). (N) Transcript levels of the ATP-generating genes *ackA* and *pgk* in FN0. (O) Box plot of the activity of the NAD biosynthesis iModulon, consisting of *nadAB* and *pnuC* that are both regulated by NadR, newly described in this study following addition of the data from this project to the *E. coli* gene expression database PRECISE1k. (P) Activity of the NAD biosynthesis iModulon in the FermNull and adapted-FermNull strains, showing a high activity of the iModulon in the engineered strain as well as attenuated activation in the adapted strains, indicating mitigated activation of NadR. All the P-values are calculated using the Welch’s t-test, with a significance threshold of 0.05.

**Methods Figure 1.**
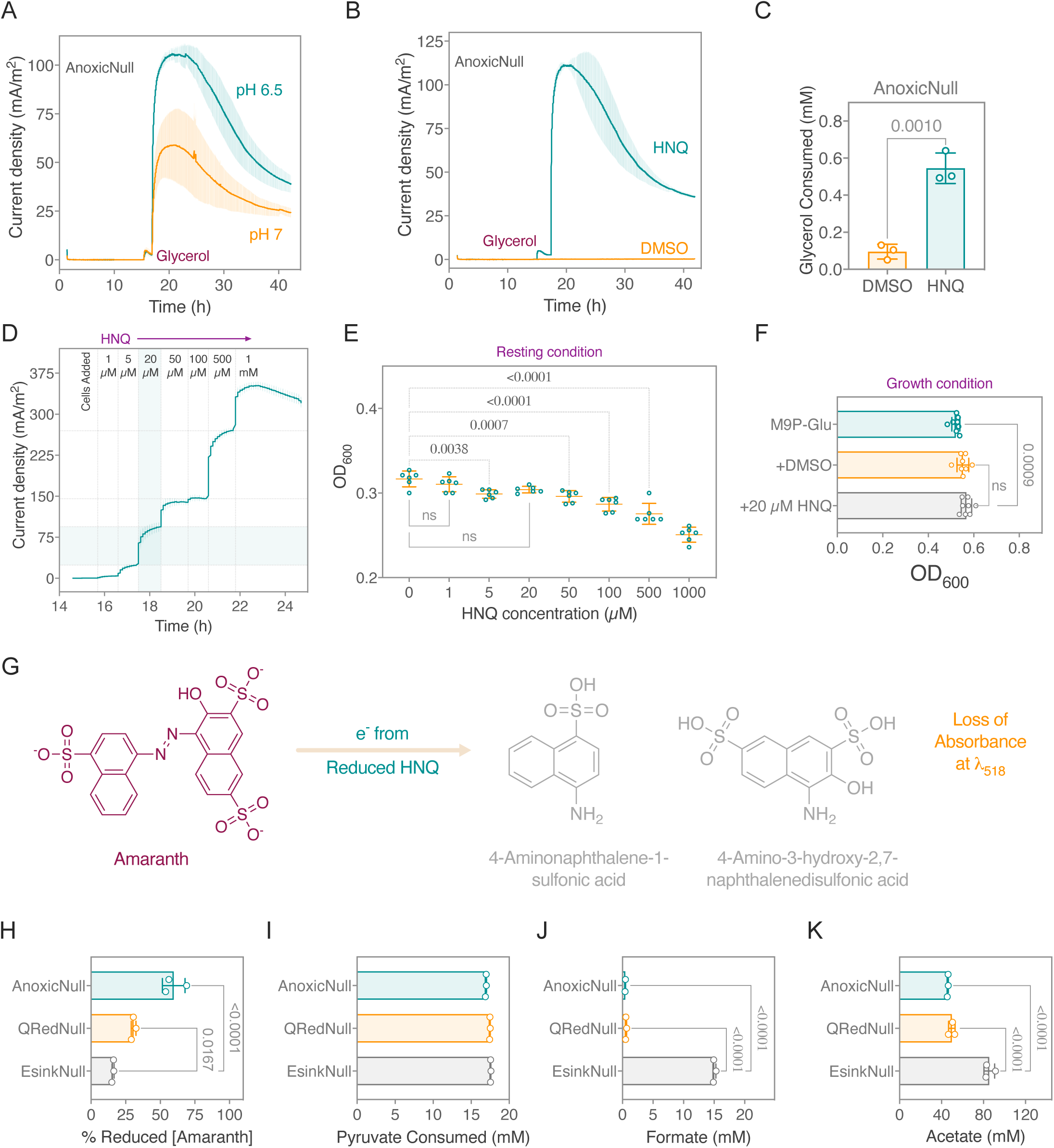
(A) Chronoamperometry of AnoxicNull at pH 6.5 and 7 under resting conditions, showing that the HNQ stimulates higher EET levels at pH 6.5 (B) Chronoamperometry of AnoxicNull with glycerol as the electron donor under resting conditions, showing that the HNQ stimulates EET whereas DMSO does not (C) Glycerol consumed by AnoxicNull in (B), showing that HNQ-mediated EET drives oxidation of glycerol. Although, AnoxicNull consumed only about 25% of the provided glycerol (2 mM). The AnoxicNull in (B) and (C) is precultured in M9P-LB, followed by biomass-generating culture in M9P-2xYT with 20 mM glycerol supplementation. The data in (A), (B), and (C) represent mean ± standard deviation obtained in triplicate BES experiments. The P-value in (C) is calculated using the Welch’s t-test, with a significance threshold of 0.05. (D) Chronoamperometry of AnoxicNull with increasing HNQ concentrations, showing that 20 µM HNQ stimulates a significant EET output. In this experiment, the cells were precultured in glycerol, and 20 mM glycerol was supplemented in the BES media. The data represents mean ± standard deviation obtained in triplicate BES experiments. (E) Cell viability assay of AnoxicNull with increasing concentrations of lawsone, showing that 20 µM HNQ does not negatively impact the viability of cells. The experiment was initiated with 1 mL of 0.5 OD_600_ cells in M9P (+2 mM glycerol) and incubated with different concentrations of HNQ for about 40 hours before taking the reported OD_600_ measurements. (F) OD_600_ of AnoxicNull in the presence of DMSO and 20 µM HNQ after 20 hours of anaerobic growth in M9P complete-growth medium with 20 mM glucose, showing that 20 µM of HNQ has no negative effect on the fermentative growth. The data represent mean ± standard deviation obtained in eight biological replicates. The P-values in (E) and (F) are calculated using one-way ANOVA, with a significance threshold of 0.05. (G) A schematic showing the structures of rose-red colored amaranth and its reduced colorless products. The degree of EET is assessed by measuring the loss of absorbance of amaranth at 518 nm. (H) The percentage reduction of amaranth by AnoxicNull, QRedNull, and EsinkNull in growth media, showing that the AnoxicNull strain reduces amaranth to the highest level, QRedNull at an intermediate level, and EsinkNull at the lowest level. This data corroborates with the EET capabilities of the strains previously elucidated from BES experiments. The data represent mean ± standard deviation obtained in three biological replicates. The P-values are calculated using one-way ANOVA. The metabolite profile of the spent BES media, showing that EsinkNull shows a similar amount of (I) pyruvate consumption yet different levels of (J) formate and (K) acetate generated, compared to AnoxicNull and QRedNull strains. This data validates the genetic backgrounds of AnoxicNull, QRedNull, and EsinkNull. The data represent mean ± standard deviation obtained in triplicate experiments. The P-values are calculated using one-way ANOVA, with a significance threshold of 0.05.

**Methods Figure 2.**
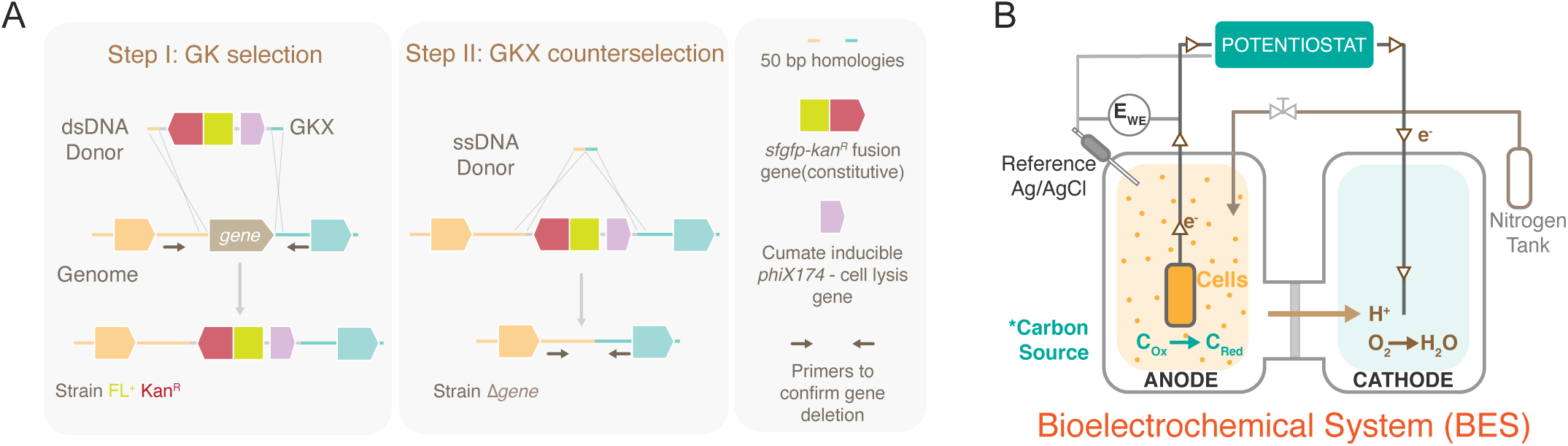
(A) A schematic showing the two-step gene deletion in BioD using the GKX selection/counterselection cassette. (B) A schematic showing the BES setup consisting of two chambers (working and counter) and three electrodes (anode, cathode, and reference).

## References

1. Zerfaß, C., Asally, M., and Soyer, O.S. (2019). Interrogating metabolism as an electron flow system. Current Opinion in Systems Biology 13, 59–67. 10.1016/j.coisb.2018.10.001.

2. Unden, G., Steinmetz, P.A., and Degreif-Dünnwald, P. (2020). The Aerobic and Anaerobic Respiratory Chain of Escherichia coli and Salmonella enterica: Enzymes and Energetics. 37.

3. Friedrich, T., and Pohl, T. (2007). NADH as Donor. EcoSal Plus 2. 10.1128/ecosalplus.3.2.4.

4. Richardson, D.J., and Cole, J.A. (2008). Respiration of Nitrate and Nitrite. EcoSal Plus 3. 10.1128/ecosal.3.2.5.

5. Gralnick, J.A., and Bond, D.R. (2023). Electron Transfer Beyond the Outer Membrane: Putting Electrons to Rest. Annual Review of Microbiology 77, 517–539. 10.1146/annurev-micro-032221-023725.

6. Jeuken, L.J.C., Hards, K., and Nakatani, Y. (2020). Extracellular Electron Transfer: Respiratory or Nutrient Homeostasis? J Bacteriol 202. 10.1128/JB.00029-20.

7. Melton, E.D., Swanner, E.D., Behrens, S., Schmidt, C., and Kappler, A. (2014). The interplay of microbially mediated and abiotic reactions in the biogeochemical Fe cycle. Nat Rev Microbiol 12, 797–808. 10.1038/nrmicro3347.

8. Shi, L., Dong, H., Reguera, G., Beyenal, H., Lu, A., Liu, J., Yu, H.-Q., and Fredrickson, J.K. (2016). Extracellular electron transfer mechanisms between microorganisms and minerals. Nat Rev Microbiol 14, 651–662. 10.1038/nrmicro.2016.93.

9. Baker, I.R., Conley, B.E., Gralnick, J.A., and Girguis, P.R. (2022). Evidence for Horizontal and Vertical Transmission of Mtr-Mediated Extracellular Electron Transfer among the Bacteria. mBio 13, e02904–21. 10.1128/mbio.02904-21.

10. Light, S.H., Su, L., Rivera-Lugo, R., Cornejo, J.A., Louie, A., Iavarone, A.T., Ajo-Franklin, C.M., and Portnoy, D.A. (2018). A flavin-based extracellular electron transfer mechanism in diverse Gram-positive bacteria. Nature 562, 140–144. 10.1038/s41586-018-0498-z.

11. Saunders, S.H., and Newman, D.K. (2018). Extracellular Electron Transfer Transcends Microbe-Mineral Interactions. Cell Host & Microbe 24, 611–613. 10.1016/j.chom.2018.10.018.

12. Logan, B.E., and Rabaey, K. (2012). Conversion of Wastes into Bioelectricity and Chemicals by Using Microbial Electrochemical Technologies. Science 337, 686–690. 10.1126/science.1217412.

13. Schievano, A., Sciarria, T.P., Vanbroekhoven, K., Wever, H.D., Puig, S., Andersen, S.J., Rabaey, K., and Pant, D. (2016). Electro-Fermentation – Merging Electrochemistry with Fermentation in Industrial Applications. Trends in Biotechnology 34, 866–878. 10.1016/j.tibtech.2016.04.007.

14. Tschirhart, T., Kim, E., McKay, R., Ueda, H., Wu, H.-C., Pottash, A.E., Zargar, A., Negrete, A., Shiloach, J., Payne, G.F., et al. (2017). Electronic control of gene expression and cell behaviour in Escherichia coli through redox signalling. Nat Commun 8, 14030. 10.1038/ncomms14030.

15. Leininger, A., Chen, J., Ramaswami, A., and Ren, Z.J. (2023). Urban circular carbon economy through electrochemically influenced microbiomes. One Earth 6, 278–289. 10.1016/j.oneear.2023.02.011.

16. Graham, A.J., Partipilo, G., Dundas, C.M., Miniel Mahfoud, I.E., Halwachs, K.N., Holwerda, A.J., Simmons, T.R., FitzSimons, T.M., Coleman, S.M., Rinehart, R., et al. (2024). Transcriptional regulation of living materials via extracellular electron transfer. Nat Chem Biol, 1–12. 10.1038/s41589-024-01628-y.

17. Gescher, J.S., Cordova, C.D., and Spormann, A.M. (2008). Dissimilatory iron reduction in Escherichia coli: identification of CymA of Shewanella oneidensis and NapC of E. coli as ferric reductases. Mol Microbiol 68, 706–719. 10.1111/j.1365-2958.2008.06183.x.

18. Goldbeck, C.P., Jensen, H.M., TerAvest, M.A., Beedle, N., Appling, Y., Hepler, M., Cambray, G., Mutalik, V., Angenent, L.T., and Ajo-Franklin, C.M. (2013). Tuning Promoter Strengths for Improved Synthesis and Function of Electron Conduits in *Escherichia coli*. ACS Synth. Biol. 2, 150–159. 10.1021/sb300119v.

19. TerAvest, M.A., and Ajo-Franklin, C.M. (2016). Transforming exoelectrogens for biotechnology using synthetic biology: Synthetic Biology of Exoelectrogens. Biotechnol. Bioeng. 113, 687–697. 10.1002/bit.25723.

20. Su, L., Fukushima, T., and Ajo-Franklin, C.M. (2020). A hybrid cyt c maturation system enhances the bioelectrical performance of engineered Escherichia coli by improving the rate-limiting step. Biosensors and Bioelectronics 165, 112312. 10.1016/j.bios.2020.112312.

21. Mouhib, M., Reggente, M., Li, L., Schuergers, N., and Boghossian, A.A. (2023). Extracellular electron transfer pathways to enhance the electroactivity of modified Escherichia coli. Joule 7, 2092–2106. 10.1016/j.joule.2023.08.006.

22. Bird, L.J., Kundu, B.B., Tschirhart, T., Corts, A.D., Su, L., Gralnick, J.A., Ajo-Franklin, C.M., and Glaven, S.M. (2021). Engineering Wired Life: Synthetic Biology for Electroactive Bacteria. ACS Synth. Biol. 10, 2808–2823. 10.1021/acssynbio.1c00335.

23. Jensen, H.M., Albers, A.E., Malley, K.R., Londer, Y.Y., Cohen, B.E., Helms, B.A., Weigele, P., Groves, J.T., and Ajo-Franklin, C.M. (2010). Engineering of a synthetic electron conduit in living cells. Proceedings of the National Academy of Sciences 107, 19213–19218. 10.1073/pnas.1009645107.

24. TerAvest, M.A., Zajdel, T.J., and Ajo-Franklin, C.M. (2014). The Mtr Pathway of *Shewanella oneidensis* MR-1 Couples Substrate Utilization to Current Production in *Escherichia coli*. CHEMELECTROCHEM 1, 1874–1879. 10.1002/celc.201402194.

25. Jensen, H.M., TerAvest, M.A., Kokish, M.G., and Ajo-Franklin, C.M. (2016). CymA and Exogenous Flavins Improve Extracellular Electron Transfer and Couple It to Cell Growth in Mtr-Expressing *Escherichia coli*. ACS Synth. Biol. 5, 679–688. 10.1021/acssynbio.5b00279.

26. Schröder, U., Nießen, J., and Scholz, F. (2003). A Generation of Microbial Fuel Cells with Current Outputs Boosted by More Than One Order of Magnitude. Angewandte Chemie International Edition 42, 2880–2883. 10.1002/anie.200350918.

27. Zhang, T., Cui, C., Chen, S., Ai, X., Yang, H., Shen, P., and Peng, Z. (2006). A novel mediatorless microbial fuel cell based on direct biocatalysis of Escherichia coli. Chemical Communications 0, 2257–2259. 10.1039/B600876C.

28. Zhang, T., Zeng, Y., Chen, S., Ai, X., and Yang, H. (2007). Improved performances of E. coli-catalyzed microbial fuel cells with composite graphite/PTFE anodes. Electrochemistry Communications 9, 349–353. 10.1016/j.elecom.2006.09.025.

29. Wang, Y.-F., Tsujimura, S., Cheng, S.-S., and Kano, K. (2007). Self-excreted mediator from Escherichia coli K-12 for electron transfer to carbon electrodes. Appl Microbiol Biotechnol 76, 1439–1446. 10.1007/s00253-007-1114-6.

30. Qiao, Y., Bao, S.-J., Li, C.M., Cui, X.-Q., Lu, Z.-S., and Guo, J. (2008). Nanostructured Polyaniline/Titanium Dioxide Composite Anode for Microbial Fuel Cells. ACS Nano 2, 113–119. 10.1021/nn700102s.

31. Qiao, Y., Li, C.M., Bao, S.-J., Lu, Z., and Hong, Y. (2008). Direct electrochemistry and electrocatalytic mechanism of evolved Escherichia coli cells in microbial fuel cells. Chem. Commun., 1290–1292. 10.1039/B719955D.

32. Xiang, K., Qiao, Y., Ching, C.B., and Li, C.M. (2009). GldA overexpressing-engineered E. coli as superior electrocatalyst for microbial fuel cells. Electrochemistry Communications 11, 1593–1595. 10.1016/j.elecom.2009.06.004.

33. Qiao, Y., Li, C.M., Lu, Z., Ling, H., Kang, A., and Chang, M.W. (2009). A time-course transcriptome analysis of Escherichia coli with direct electrochemistry behavior in microbial fuel cells. Chem. Commun., 6183–6185. 10.1039/B912003C.

34. Liu, J., Yong, Y.-C., Song, H., and Li, C.M. (2012). Activation Enhancement of Citric Acid Cycle to Promote Bioelectrocatalytic Activity of *arcA* Knockout *Escherichia coli* Toward High-Performance Microbial Fuel Cell. ACS Catal. 2, 1749–1752. 10.1021/cs3003808.

35. Yong, Y.-C., Yu, Y.-Y., Yang, Y., Li, C.M., Jiang, R., Wang, X., Wang, J.-Y., and Song, H. (2012). Increasing intracellular releasable electrons dramatically enhances bioelectricity output in microbial fuel cells. Electrochemistry Communications 19, 13–16. 10.1016/j.elecom.2012.03.002.

36. Logan, B.E., Rossi, R., Ragab, A., and Saikaly, P.E. (2019). Electroactive microorganisms in bioelectrochemical systems. Nat Rev Microbiol 17, 307–319. 10.1038/s41579-019-0173-x.

37. Glasser, N.R., Wang, B.X., Hoy, J.A., and Newman, D.K. (2017). The Pyruvate and α-Ketoglutarate Dehydrogenase Complexes of Pseudomonas aeruginosa Catalyze Pyocyanin and Phenazine-1-carboxylic Acid Reduction via the Subunit Dihydrolipoamide Dehydrogenase*. Journal of Biological Chemistry 292, 5593–5607. 10.1074/jbc.M116.772848.

38. Ciemniecki, J.A., and Newman, D.K. (2023). NADH dehydrogenases are the predominant phenazine reductases in the electron transport chain of Pseudomonas aeruginosa. Molecular Microbiology 119, 560–573. 10.1111/mmi.15049.

39. Hinks, J., Han, E.J.Y., Wang, V.B., Seviour, T.W., Marsili, E., Loo, J.S.C., and Wuertz, S. (2016). Naphthoquinone glycosides for bioelectroanalytical enumeration of the faecal indicator Escherichia coli. Microbial Biotechnology 9, 746–757. 10.1111/1751-7915.12373.

40. Egbert, R.G., Rishi, H.S., Adler, B.A., McCormick, D.M., Toro, E., Gill, R.T., and Arkin, A.P. (2019). A versatile platform strain for high-fidelity multiplex genome editing. Nucleic Acids Research 47, 3244–3256. 10.1093/nar/gkz085.

41. Dharmadi, Y., Murarka, A., and Gonzalez, R. (2006). Anaerobic fermentation of glycerol by Escherichia coli: A new platform for metabolic engineering. Biotechnology and Bioengineering 94, 821–829. 10.1002/bit.21025.

42. Clomburg, J.M., Cintolesi, A., and Gonzalez, R. (2022). In silico and in vivo analyses reveal key metabolic pathways enabling the fermentative utilization of glycerol in Escherichia coli. Microbial Biotechnology 15, 289–304. 10.1111/1751-7915.13938.

43. Koland, J.G., Miller, M.J., and Gennis, R.B. (1984). Reconstitution of the membrane-bound, ubiquinone-dependent pyruvate oxidase respiratory chain of Escherichia coli with the cytochrome d terminal oxidase. Biochemistry 23, 445–453. 10.1021/bi00298a008.

44. Rychel, K., Decker, K., Sastry, A.V., Phaneuf, P.V., Poudel, S., and Palsson, B.O. (2021). iModulonDB: a knowledgebase of microbial transcriptional regulation derived from machine learning. Nucleic Acids Research 49, D112–D120. 10.1093/nar/gkaa810.

45. Lamoureux, C.R., Decker, K.T., Sastry, A.V., Rychel, K., Gao, Y., McConn, J.L., Zielinski, D.C., and Palsson, B.O. (2023). A multi-scale expression and regulation knowledge base for Escherichia coli. Nucleic Acids Research 51, 10176–10193. 10.1093/nar/gkad750.

46. Rothery, R.A., Chatterjee, I., Kiema, G., McDermott, M.T., and Weiner, J.H. (1998). Hydroxylated naphthoquinones as substrates for Escherichia coli anaerobic reductases. Biochemical Journal 332, 35–41. 10.1042/bj3320035.

47. Tellurite reductase activity of nitrate reductase is responsible for the basal resistance of Escherichia coli to tellurite - PubMed https://pubmed.ncbi.nlm.nih.gov/9141681/.

48. Harrington, T.D., Tran, V.N., Mohamed, A., Renslow, R., Biria, S., Orfe, L., Call, D.R., and Beyenal, H. (2015). The mechanism of neutral red-mediated microbial electrosynthesis in Escherichia coli: menaquinone reduction. Bioresource Technology 192, 689–695. 10.1016/j.biortech.2015.06.037.

49. Tolar, J.G., Li, S., and Ajo-Franklin, C.M. (2023). The Differing Roles of Flavins and Quinones in Extracellular Electron Transfer in Lactiplantibacillus plantarum. Appl Environ Microbiol 89, e01313–22. 10.1128/aem.01313-22.

50. Li, S., De Groote Tavares, C., Tolar, J.G., and Ajo-Franklin, C.M. (2024). Selective bioelectronic sensing of pharmacologically relevant quinones using extracellular electron transfer in *Lactiplantibacillus plantarum*. Biosensors and Bioelectronics 243, 115762. 10.1016/j.bios.2023.115762.

51. Imlay, J., and Fridovich, I. Exogenous Quinones Directly Inhibit the Respiratory NADH Dehydrogenase in Escherichia co/i’. 10.

52. Dancey, G.F., and Shapiro, B.M. (1976). The NADH dehydrogenase of the respiratory chain of Escherichia coli. II. Kinetics of the purified enzyme and the effects of antibodies elicited against it on membrane-bound and free enzyme. J Biol Chem 251, 5921–5928.

53. Hayashi, M., Hasegawa, K., Oguni, Y., and Unemoto, T. (1990). Characterization of FMN-dependent NADH-quinone reductase induced by menadione in Escherichia coli. Biochimica et Biophysica Acta (BBA) - General Subjects 1035, 230–236. 10.1016/0304-4165(90)90122-D.

54. Unemoto, T., Shimada, H., and Hayashi, M. (1992). Chemical structures critical for the induction of FMN-dependent NADH-quinone reductase in Escherichia coli. Biochim Biophys Acta 1099, 170–174.

55. Promoted performance of microbial fuel cells using Escherichia coli cells with multiple-knockout of central metabolism genes.

56. Sawers, R.G., and Clark, D.P. (2020). Fermentative Pyruvate and Acetyl-Coenzyme A Metabolism. 38.

57. Sánchez, L.B., Elmendorf, H., Nash, T.E., and Müller, M. 2001 NAD(P)H:menadione oxidoreductase of the amitochondriate eukaryote Giardia lamblia: a simpler homologue of the vertebrate enzymeThe GenBank accession number for the glQR1-encoding sequence reported in this paper is AF321405. Microbiology 147, 561–570. 10.1099/00221287-147-3-561.

58. Sandoval, J.M., Arenas, F.A., and Vásquez, C.C. (2011). Glucose-6-Phosphate Dehydrogenase Protects Escherichia coli from Tellurite-Mediated Oxidative Stress. PLOS ONE 6, e25573. 10.1371/journal.pone.0025573.

59. Greenberg, J.T., and Demple, B. (1989). A global response induced in Escherichia coli by redox-cycling agents overlaps with that induced by peroxide stress. J Bacteriol 171, 3933–3939. 10.1128/jb.171.7.3933-3939.1989.

60. Koch, K., Strandback, E., Jha, S., Richter, G., Bourgeois, B., Madl, T., and Macheroux, P. (2019). Oxidative stress-induced structural changes in the microtubule-associated flavoenzyme Irc15p from Saccharomyces cerevisiae. Protein Sci 28, 176–190. 10.1002/pro.3517.

61. Smirnova, G.V., Muzyka, N.G., Glukhovchenko, M.N., and Oktyabrsky, O.N. (2000). Effects of menadione and hydrogen peroxide on glutathione status in growing Escherichia coli. Free Radical Biology and Medicine 28, 1009–1016. 10.1016/S0891-5849(99)00256-7.

62. Mauzeroll, J., Bard, A.J., Owhadian, O., and Monks, T.J. (2004). Menadione metabolism to thiodione in hepatoblastoma by scanning electrochemical microscopy. Proceedings of the National Academy of Sciences 101, 17582–17587. 10.1073/pnas.0407613101.

63. McMillan, D.C., Sarvate, S.D., Oatis, J.E., and Jollow, D.J. (2004). Role of oxidant stress in lawsone-induced hemolytic anemia. Toxicol Sci 82, 647–655. 10.1093/toxsci/kfh288.

64. Castro, F.A.V., Herdeiro, R.S., Panek, A.D., Eleutherio, E.C.A., and Pereira, M.D. (2007). Menadione stress in Saccharomyces cerevisiae strains deficient in the glutathione transferases. Biochimica et Biophysica Acta (BBA) - General Subjects 1770, 213–220. 10.1016/j.bbagen.2006.10.013.

65. Green, A.R., Hayes, R.P., Xun, L., and Kang, C. (2012). Structural understanding of the glutathione-dependent reduction mechanism of glutathionyl-hydroquinone reductases. J Biol Chem 287, 35838–35848. 10.1074/jbc.M112.395541.

66. Belorgey, D., Lanfranchi, D.A., and Davioud-Charvet, E. (2013). 1,4-Naphthoquinones and Others NADPH-Dependent Glutathione Reductase-Catalyzed Redox Cyclers as Antimalarial Agents. Curr Pharm Des 19, 2512–2528.

67. Loi, V.V., Rossius, M., and Antelmann, H. (2015). Redox regulation by reversible protein S-thiolation in bacteria. Frontiers in Microbiology 6.

68. Liebeke, M., Pöther, D.-C., Van Duy, N., Albrecht, D., Becher, D., Hochgräfe, F., Lalk, M., Hecker, M., and Antelmann, H. (2008). Depletion of thiol-containing proteins in response to quinones in Bacillus subtilis. Molecular Microbiology 69, 1513–1529. 10.1111/j.1365-2958.2008.06382.x.

69. Sun, S., Zhang, Y., Xu, W., Yang, R., Yang, Y., Guo, J., Ma, Q., Ma, K., Zhang, J., and Xu, J. (2022). Plumbagin reduction by thioredoxin reductase 1 possesses synergy effects with GLUT1 inhibitor on KEAP1-mutant NSCLC cells. Biomedicine & Pharmacotherapy 146, 112546. 10.1016/j.biopha.2021.112546.

70. Wang, P.F., Marcinkeviciene, J., Williams, C.H., and Blanchard, J.S. (1998). Thioredoxin reductase-thioredoxin fusion enzyme from Mycobacterium leprae: comparison with the separately expressed thioredoxin reductase. Biochemistry 37, 16378–16389. 10.1021/bi980754e.

71. Nickerson, W.J., Falcone, G., and Strauss, G. (1963). Studies on Quinone-Thioethers. I. Mechanism of Formation and Properties of Thiodione*. Biochemistry 2, 537–543. 10.1021/bi00903a025.

72. Liu, G., Zhou, J., Fu, Q.S., and Wang, J. (2009). The Escherichia coli Azoreductase AzoR Is Involved in Resistance to Thiol-Specific Stress Caused by Electrophilic Quinones. J Bacteriol 191, 6394–6400. 10.1128/JB.00552-09.

73. Liu, G., Zhou, J., Wang, J., Zhou, M., Lu, H., and Jin, R. (2009). Acceleration of azo dye decolorization by using quinone reductase activity of azoreductase and quinone redox mediator. Bioresource Technology 100, 2791–2795. 10.1016/j.biortech.2008.12.040.

74. Valiauga, B., Williams, E.M., Ackerley, D.F., and Čėnas, N. (2017). Reduction of quinones and nitroaromatic compounds by Escherichia coli nitroreductase A (NfsA): Characterization of kinetics and substrate specificity. Archives of Biochemistry and Biophysics 614, 14–22. 10.1016/j.abb.2016.12.005.

75. Zenno, S., Koike, H., Tanokura, M., and Saigo, K. (1996). Conversion of NfsB, a minor Escherichia coli nitroreductase, to a flavin reductase similar in biochemical properties to FRase I, the major flavin reductase in Vibrio fischeri, by a single amino acid substitution. Journal of Bacteriology 178, 4731–4733. 10.1128/jb.178.15.4731-4733.1996.

76. Adams, M.A., and Jia, Z. (2006). Modulator of Drug Activity B from Escherichia coli: Crystal Structure of a Prokaryotic Homologue of DT-diaphorase. Journal of Molecular Biology 359, 455–465. 10.1016/j.jmb.2006.03.053.

77. Adams, M.A., and Jia, Z. (2005). Structural and Biochemical Evidence for an Enzymatic Quinone Redox Cycle in Escherichia coli: IDENTIFICATION OF A NOVEL QUINOL MONOOXYGENASE * [boxs]. Journal of Biological Chemistry 280, 8358–8363. 10.1074/jbc.M412637200.

78. Patridge, E.V., and Ferry, J.G. (2006). WrbA from Escherichia coli and Archaeoglobus fulgidus Is an NAD(P)H:Quinone Oxidoreductase. Journal of Bacteriology 188, 3498–3506. 10.1128/jb.188.10.3498-3506.2006.

79. Gonzalez, C.F., Ackerley, D.F., Lynch, S.V., and Matin, A. (2005). ChrR, a Soluble Quinone Reductase of Pseudomonas putida That Defends against H2O2*. Journal of Biological Chemistry 280, 22590–22595. 10.1074/jbc.M501654200.

80. Lyngberg, L., Healy, J., Bartlett, W., Miller, S., Conway, S.J., Booth, I.R., and Rasmussen, T. (2011). KefF, the Regulatory Subunit of the Potassium Efflux System KefC, Shows Quinone Oxidoreductase Activity. Journal of Bacteriology 193, 4925–4932. 10.1128/jb.05272-11.

81. Healy, J., Ekkerman, S., Pliotas, C., Richard, M., Bartlett, W., Grayer, S.C., Morris, G.M., Miller, S., Booth, I.R., Conway, S.J., et al. (2014). Understanding the Structural Requirements for Activators of the Kef Bacterial Potassium Efflux System. Biochemistry 53, 1982–1992. 10.1021/bi5001118.

82. Lee, C., Shin, J., and Park, C. (2013). Novel regulatory system nemRA–gloA for electrophile reduction in Escherichia coli K-12. Molecular Microbiology 88, 395–412. 10.1111/mmi.12192.

83. Iwadate, Y., Honda, H., Sato, H., Hashimoto, M., and Kato, J. (2011). Oxidative stress sensitivity of engineered Escherichia coli cells with a reduced genome. FEMS Microbiology Letters 322, 25–33. 10.1111/j.1574-6968.2011.02331.x.

84. Ostrowski, J., Barber, M.J., Rueger, D.C., Miller, B.E., Siegel, L.M., and Kredich, N.M. (1989). Characterization of the Flavoprotein Moieties of NADPH-Sulfite Reductase from Salmonella typhimurium and Escherichia coli. Journal of Biological Chemistry 264, 15796–15808. 10.1016/S0021-9258(18)71547-0.

85. Arenas, F.A., Díaz, W.A., Leal, C.A., Pérez-Donoso, J.M., Imlay, J.A., and Vásquez, C.C. (2010). The Escherichia coli btuE gene, encodes a glutathione peroxidase that is induced under oxidative stress conditions. Biochemical and Biophysical Research Communications 398, 690–694. 10.1016/j.bbrc.2010.07.002.

86. Sheppard, C.A., Trimmer, E.E., and Matthews, R.G. (1999). Purification and Properties of NADH-Dependent 5,10-Methylenetetrahydrofolate Reductase (MetF) fromEscherichia coli. Journal of Bacteriology 181, 718–725. 10.1128/jb.181.3.718-725.1999.

87. Pérez, J.M., Arenas, F.A., Pradenas, G.A., Sandoval, J.M., and Vásquez, C.C. (2008). Escherichia coli YqhD Exhibits Aldehyde Reductase Activity and Protects from the Harmful Effect of Lipid Peroxidation-derived Aldehydes*. Journal of Biological Chemistry 283, 7346–7353. 10.1074/jbc.M708846200.

88. Misal, S.A., and Gawai, K.R. (2018). Azoreductase: a key player of xenobiotic metabolism. Bioresources and Bioprocessing 5, 17. 10.1186/s40643-018-0206-8.

89. Henshke, Y., Shemer, B., and Belkin, S. (2021). The Escherichia coli azoR gene promoter: A new sensing element for microbial biodetection of trace explosives. Current Research in Biotechnology 3, 21–28. 10.1016/j.crbiot.2021.01.003.

90. Rau, J., and Stolz, A. (2003). Oxygen-Insensitive Nitroreductases NfsA and NfsB of Escherichia coli Function under Anaerobic Conditions as Lawsone-Dependent Azo Reductases. Applied and Environmental Microbiology 69, 3448–3455. 10.1128/AEM.69.6.3448-3455.2003.

91. Zenno, S., Koike, H., Kumar, A.N., Jayaraman, R., Tanokura, M., and Saigo, K. (1996). Biochemical characterization of NfsA, the Escherichia coli major nitroreductase exhibiting a high amino acid sequence homology to Frp, a Vibrio harveyi flavin oxidoreductase. J Bacteriol 178, 4508–4514. 10.1128/jb.178.15.4508-4514.1996.

92. Bryant, D.W., McCalla, D.R., Leeksma, M., and Laneuville, P. (1981). Type I nitroreductases of Escherichia coli. Can J Microbiol 27, 81–86. 10.1139/m81-013.

93. Pinske, C., and Sawers, R.G. (2016). Anaerobic Formate and Hydrogen Metabolism. EcoSal Plus 7. 10.1128/ecosalplus.ESP-0011-2016.

94. Dalldorf, C., Rychel, K., Szubin, R., Hefner, Y., Patel, A., Zielinski, D., and Palsson, B. (2023). The hallmarks of a tradeoff in transcriptomes that balances stress and growth functions. Preprint, 10.21203/rs.3.rs-2729651/v1.

95. Orth, J.D., Thiele, I., and Palsson, B.Ø. (2010). What is flux balance analysis? Nat Biotechnol 28, 245–248. 10.1038/nbt.1614.

96. Monk, J.M., Lloyd, C.J., Brunk, E., Mih, N., Sastry, A., King, Z., Takeuchi, R., Nomura, W., Zhang, Z., Mori, H., et al. (2017). iML1515, a knowledgebase that computes Escherichia coli traits. Nat Biotechnol 35, 904–908. 10.1038/nbt.3956.

97. O’Brien, E.J., Lerman, J.A., Chang, R.L., Hyduke, D.R., and Palsson, B.Ø. (2013). Genome-scale models of metabolism and gene expression extend and refine growth phenotype prediction. Mol Syst Biol 9, 693. 10.1038/msb.2013.52.

98. Lloyd, C.J., Ebrahim, A., Yang, L., King, Z.A., Catoiu, E., O’Brien, E.J., Liu, J.K., and Palsson, B.O. (2018). COBRAme: A computational framework for genome-scale models of metabolism and gene expression. PLoS Comput Biol 14, e1006302. 10.1371/journal.pcbi.1006302.

99. Zhao, J., Li, F., Cao, Y., Zhang, X., Chen, T., Song, H., and Wang, Z. (2021). Microbial extracellular electron transfer and strategies for engineering electroactive microorganisms. Biotechnology Advances 53, 107682. 10.1016/j.biotechadv.2020.107682.

100. Gemünde, A., Lai, B., Pause, L., Krömer, J., and Holtmann, D. Redox Mediators in Microbial Electrochemical Systems. ChemElectroChem n/a, e202200216. 10.1002/celc.202200216.

101. Price-Whelan, A., Dietrich, L.E.P., and Newman, D.K. (2007). Pyocyanin Alters Redox Homeostasis and Carbon Flux through Central Metabolic Pathways in Pseudomonas aeruginosa PA14. Journal of Bacteriology 189, 6372–6381. 10.1128/jb.00505-07.

102. Glasser, N.R., Kern, S.E., and Newman, D.K. (2014). Phenazine redox cycling enhances anaerobic survival in Pseudomonas aeruginosa by facilitating generation of ATP and a proton-motive force. Molecular Microbiology 92, 399–412. 10.1111/mmi.12566.

103. Jo, J., Price-Whelan, A., Cornell, W.C., and Dietrich, L.E.P. (2020). Interdependency of Respiratory Metabolism and Phenazine-Associated Physiology in Pseudomonas aeruginosa PA14. Journal of Bacteriology 202, 10.1128/jb.00700-19. 10.1128/jb.00700-19.

104. Schulz-Mirbach, H., Dronsella, B., He, H., and Erb, T.J. (2023). Creating new-to-nature carbon fixation: A guide. Metabolic Engineering. 10.1016/j.ymben.2023.12.012.

105. Biner, O., Fedor, J.G., Yin, Z., and Hirst, J. (2020). Bottom-Up Construction of a Minimal System for Cellular Respiration and Energy Regeneration. ACS Synth. Biol. 9, 1450–1459. 10.1021/acssynbio.0c00110.

106. Bergkessel, M., Basta, D.W., and Newman, D.K. (2016). The physiology of growth arrest: uniting molecular and environmental microbiology. Nat Rev Microbiol 14, 549–562. 10.1038/nrmicro.2016.107.

107. Lovley, D.R., and Coates, J.D. (2000). Novel forms of anaerobic respiration of environmental relevance. Current Opinion in Microbiology 3, 252–256. 10.1016/S1369-5274(00)00085-0.

108. Wolf, M., Kappler, A., Jiang, J., and Meckenstock, R.U. (2009). Effects of Humic Substances and Quinones at Low Concentrations on Ferrihydrite Reduction by Geobacter metallireducens. Environ. Sci. Technol. 43, 5679–5685. 10.1021/es803647r.

109. Wang, X., Liu, G., Zhou, J., Wang, J., Jin, R., and Lv, H. (2011). Quinone-mediated reduction of selenite and tellurite by Escherichia coli. Bioresour Technol 102, 3268–3271. 10.1016/j.biortech.2010.11.078.

110. Borghese, R., Baccolini, C., Francia, F., Sabatino, P., Turner, R.J., and Zannoni, D. (2014). Reduction of chalcogen oxyanions and generation of nanoprecipitates by the photosynthetic bacterium Rhodobacter capsulatus. Journal of Hazardous Materials 269, 24–30. 10.1016/j.jhazmat.2013.12.028.

111. Russ, R., Rau, J., and Stolz, A. (2000). The Function of Cytoplasmic Flavin Reductases in the Reduction of Azo Dyes by Bacteria. Applied and Environmental Microbiology 66, 1429–1434. 10.1128/AEM.66.4.1429-1434.2000.

112. Rau, J., Knackmuss, H.-J., and Stolz, A. (2002). Effects of Different Quinoid Redox Mediators on the Anaerobic Reduction of Azo Dyes by Bacteria. Environ. Sci. Technol. 36, 1497–1504. 10.1021/es010227+.

113. Lovley, D.R., Coates, J.D., Blunt-Harris, E.L., Phillips, E.J.P., and Woodward, J.C. (1996). Humic substances as electron acceptors for microbial respiration. Nature 382, 445–448. 10.1038/382445a0.

114. Newman, D.K., and Kolter, R. (2000). A role for excreted quinones in extracellular electron transfer. Nature 405, 94–97. 10.1038/35011098.

115. Mevers, E., Su, L., Pishchany, G., Baruch, M., Cornejo, J., Hobert, E., Dimise, E., Ajo-Franklin, C.M., and Clardy, J. (2019). An elusive electron shuttle from a facultative anaerobe. eLife 8, e48054. 10.7554/eLife.48054.

116. Yamazaki, S., Kaneko, T., Taketomo, N., Kano, K., and Ikeda, T. (2002). Glucose Metabolism of Lactic Acid Bacteria Changed by Quinone-mediated Extracellular Electron Transfer. Bioscience, Biotechnology, and Biochemistry 66, 2100–2106. 10.1271/bbb.66.2100.

117. Li, X., Liu, L., Liu, T., Yuan, T., Zhang, W., Li, F., Zhou, S., and Li, Y. (2013). Electron transfer capacity dependence of quinone-mediated Fe(III) reduction and current generation by *Klebsiella pneumoniae* L17. Chemosphere 92, 218–224. 10.1016/j.chemosphere.2013.01.098.

118. Moyo, C.E., Beckett, R.P., Trifonova, T.V., and Minibayeva, F.V. (2017). Extracellular redox cycling and hydroxyl radical production occurs widely in lichenized Ascomycetes. Fungal Biology 121, 582–588. 10.1016/j.funbio.2017.03.005.

119. Turick, C.E. The physiological role and characterization of melanin produced by Shewanella algae BrY. 99.

120. Turick, C.E., Tisa, L.S., and Caccavo, Jr., F. (2002). Melanin Production and Use as a Soluble Electron Shuttle for Fe(III) Oxide Reduction and as a Terminal Electron Acceptor by Shewanella algae BrY. Appl Environ Microbiol 68, 2436–2444. 10.1128/AEM.68.5.2436-2444.2002.

121. Meganathan, R., and Kwon, O. (2009). Biosynthesis of Menaquinone (Vitamin K2) and Ubiquinone (Coenzyme Q). EcoSal Plus 3, 10.1128/ecosalplus.3.6.3.3. 10.1128/ecosalplus.3.6.3.3.

122. Daisley, B.A., Koenig, D., Engelbrecht, K., Doney, L., Hards, K., Al, K.F., Reid, G., and Burton, J.P. (2021). Emerging connections between gut microbiome bioenergetics and chronic metabolic diseases. Cell Reports 37, 110087. 10.1016/j.celrep.2021.110087.

123. Stevens, E.T., Van Beeck, W., Blackburn, B., Tejedor-Sanz, S., Rasmussen, A.R.M., Carter, M.E., Mevers, E., Ajo-Franklin, C.M., and Marco, M.L. (2023). *Lactiplantibacillus plantarum* uses ecologically relevant, exogenous quinones for extracellular electron transfer. mBio 14, e02234–23. 10.1128/mbio.02234-23.

124. Widhalm, J.R., and Rhodes, D. (2016). Biosynthesis and molecular actions of specialized 1,4-naphthoquinone natural products produced by horticultural plants. Hortic Res 3, 16046. 10.1038/hortres.2016.46.

125. Meyer, G.W., Bahamon Naranjo, M.A., and Widhalm, J.R. (2021). Convergent evolution of plant specialized 1,4-naphthoquinones: metabolism, trafficking, and resistance to their allelopathic effects. Journal of Experimental Botany 72, 167–176. 10.1093/jxb/eraa462.

126. Clomburg, J.M., and Gonzalez, R. (2013). Anaerobic fermentation of glycerol: a platform for renewable fuels and chemicals. Trends in Biotechnology 31, 20–28. 10.1016/j.tibtech.2012.10.006.

127. Deatherage, D.E., and Barrick, J.E. (2014). Identification of Mutations in Laboratory-Evolved Microbes from Next-Generation Sequencing Data Using breseq. In Engineering and Analyzing Multicellular Systems: Methods and Protocols, L. Sun and W. Shou, eds. (Springer), pp. 165–188. 10.1007/978-1-4939-0554-6_12.

128. Seemann, T. (2014). Prokka: rapid prokaryotic genome annotation. Bioinformatics 30, 2068–2069. 10.1093/bioinformatics/btu153.

129. Bankevich, A., Nurk, S., Antipov, D., Gurevich, A.A., Dvorkin, M., Kulikov, A.S., Lesin, V.M., Nikolenko, S.I., Pham, S., Prjibelski, A.D., et al. (2012). SPAdes: A New Genome Assembly Algorithm and Its Applications to Single-Cell Sequencing. Journal of Computational Biology 19, 455–477. 10.1089/cmb.2012.0021.

130. Reis, A.C., and Salis, H.M. (2020). An Automated Model Test System for Systematic Development and Improvement of Gene Expression Models. ACS Synth. Biol. 9, 3145–3156. 10.1021/acssynbio.0c00394.

131. Hossain, A., Lopez, E., Halper, S.M., Cetnar, D.P., Reis, A.C., Strickland, D., Klavins, E., and Salis, H.M. (2020). Automated design of thousands of nonrepetitive parts for engineering stable genetic systems. Nat Biotechnol 38, 1466–1475. 10.1038/s41587-020-0584-2.

132. Park, Y., Espah Borujeni, A., Gorochowski, T.E., Shin, J., and Voigt, C.A. (2020). Precision design of stable genetic circuits carried in highly-insulated E. coli genomic landing pads. Molecular Systems Biology 16, e9584. 10.15252/msb.20209584.

133. Meyer, A.J., Segall-Shapiro, T.H., Glassey, E., Zhang, J., and Voigt, C.A. (2019). Escherichia coli “Marionette” strains with 12 highly optimized small-molecule sensors. Nat Chem Biol 15, 196–204. 10.1038/s41589-018-0168-3.

134. Monk, J.M., Koza, A., Campodonico, M.A., Machado, D., Seoane, J.M., Palsson, B.O., Herrgård, M.J., and Feist, A.M. (2016). Multi-omics Quantification of Species Variation of Escherichia coli Links Molecular Features with Strain Phenotypes. Cell Systems 3, 238–251.e12. 10.1016/j.cels.2016.08.013.

135. Heirendt, L., Arreckx, S., Pfau, T., Mendoza, S.N., Richelle, A., Heinken, A., Haraldsdóttir, H.S., Wachowiak, J., Keating, S.M., Vlasov, V., et al. (2019). Creation and analysis of biochemical constraint-based models using the COBRA Toolbox v.3.0. Nat Protoc 14, 639–702. 10.1038/s41596-018-0098-2.

136. Patel, A., McGrosso, D., Hefner, Y., Campeau, A., Sastry, A.V., Maurya, S., Rychel, K., Gonzalez, D.J., and Palsson, B.O. (2023). Proteome allocation is linked to transcriptional regulation through a modularized transcriptome. Preprint at bioRxiv, 10.1101/2023.02.20.529291.

137. Sastry, A.V., Poudel, S., Rychel, K., Yoo, R., Lamoureux, C.R., Chauhan, S., Haiman, Z.B., Bulushi, T.A., Seif, Y., and Palsson, B.O. (2021). Mining all publicly available expression data to compute dynamic microbial transcriptional regulatory networks. Preprint at bioRxiv, 10.1101/2021.07.01.450581.

